# Cortical neurons accumulate point mutations at similar rates in mice and humans

**DOI:** 10.64898/2026.11.21.768276

**Authors:** Matin Saeidi, Juyoun Yoo, Anastasia Stolyarova, Franck Polleux, Molly Przeworski, Natanael Spisak

**Author notes:** These authors co-supervised this work (Molly Przeworski, Natanael Spisak).

## Abstract

Across vertebrate species, it has long been known that germline mutation rates per year are inversely proportional to reproductive ages. A similar pattern was recently reported in mammals for mutations in colonic epithelium, leading to speculation that this inter-species scaling may apply to many cell types. We tested whether a similar relationship holds in postmitotic cells, by estimating point mutation rates in mouse cortical neurons from PacBio HiFi circular consensus sequencing data. Our analysis revealed a significant effect of age on the number of point mutations, of 25.9 mutations per diploid genome per year (95% CI: [17.7, 34.1]). This rate is comparable to the one reported for human neurons, indicating that somatic rates need not scale with lifespan. Our findings further demonstrate that a readily available sequencing method can be used to reliably identify somatic mutations at single-cell resolution.

## Introduction

Mutation rate differences among species are informative about sources of mutations and the selection pressures that shape their evolution. In the germline, it has long been observed that vertebrates with longer generation times have lower point mutation rates per year ^1–4^. An equivalent observation, confirmed in pedigree data, is that vertebrate species tend to have very similar germline mutation rates at their typical ages of reproduction (reviewed in de Manuel et al.^5^ ). Explanations for this phenomenon usually invoke strong purifying selection against germline mutations, which are constitutive in offspring ^6,7^. Given that most germline point mutations appear to be triggered by damage, rather than errors during whole genome replication cycles^8^, the observation of decreased mutation rates per year in long-lived species further suggests that their damage rates are lower or repair efficiencies/accuracies higher.

Recently, a study found a similar scaling across mammalian species in colonic crypts. In laser-microdissected samples, the somatic point mutation rate per year was inversely proportional to lifespan or equivalently, the burden of somatic mutation was similar at the end of lifespan ^9^. As one example, mouse colonic stem cells accumulated about 800 mutations per year compared to about 50 in humans, a *∼*17-fold difference, on par with the *∼*23-fold difference in lifespan. Given the study design, most of the mutations that were detected likely arose along the lineage of the stem cells that give rise to and maintain a crypt, rather than in its differentiated cell progeny. The authors suggest that the lower rates in longer-lived mammals reflect strong selection on somatic mutations, potentially because they contribute to aging processes^9^. In colon, as in the germline, the spectrum of point mutations is broadly similar across species, further indicating that similar mutagenic processes are at work^9, 10^.

On first principles, we might expect selection pressures on somatic cell types to depend on a number of features, including their contribution to organismal fitness, how long-lived they are, as well as tissue architecture^5,11,12^. Little is known empirically, however, other than for mouse blood stem cells, in which the mutation rate was estimated to be only 3-fold higher than in humans ^13^. Neurons are of particular interest in this regard, given that they are postmitotic cells and need to be structurally and functionally maintained throughout the lifespan of the animal—many decades in humans but only a couple of years in mice. In humans, neurons accumulate 16 to 23 mutations per year ^14–17^, due to the misrepair of DNA damage^18^. In mice, a recent study of the effect of caloric restriction on mutation rates indicated that in the cerebellum, the number of point mutations that accumulated in 19 months is similar to what is predicted for humans of the same age ^19^. Here, we focus on mouse cortical neurons, in order to characterize how mutations accumulate with age in a postmitotic cell population.

## Results and discussion

To date, the study of DNA mutations at single-cell resolution has required laborious experimental approaches, including clonal expansions^20–23^, whole-genome amplification of single cells^14,16^, or duplex short-read sequencing ^15^. More recently, it has become possible to take a simpler approach, based on an off-the-shelf technology: PacBio circular consensus sequencing (CCS) ^24^. CCS libraries are straightforward to prepare and involve no amplification, which can introduce mismatches and other artifacts. This approach has been used to call non-crossover events in primate sperm and testes samples^25–27^ and mutations in tissue or organismal samples collected for the purposes of genome assembly ^28^. We apply it here with minor modifications inspired by the HiDEF-seq protocol^29^, in order to decrease errors relative to the number of mutations carried by a single molecule. We evaluate the reliability of our approach after modeling residual errors and filtering them out in downstream analysis.

### Bioinformatic pipeline and error rates

#### Study design

To call somatic mutations, we sequenced cortical neuron nuclei isolated from seven mice aged 12 to 72 weeks. Because neurons only constitute *∼*50% of all cells contained in the adult mouse cortex, we isolated nuclei from neurons using fluorescence activated sorting (FACS) of NeuN-positive nuclei^29^ (see Methods). We used circular consensus sequencing of one library from each of the seven isogenic C57BL/6J mice and two libraries from the 60-week-old animal (Fig. 1, see Methods).

**Figure 1.**
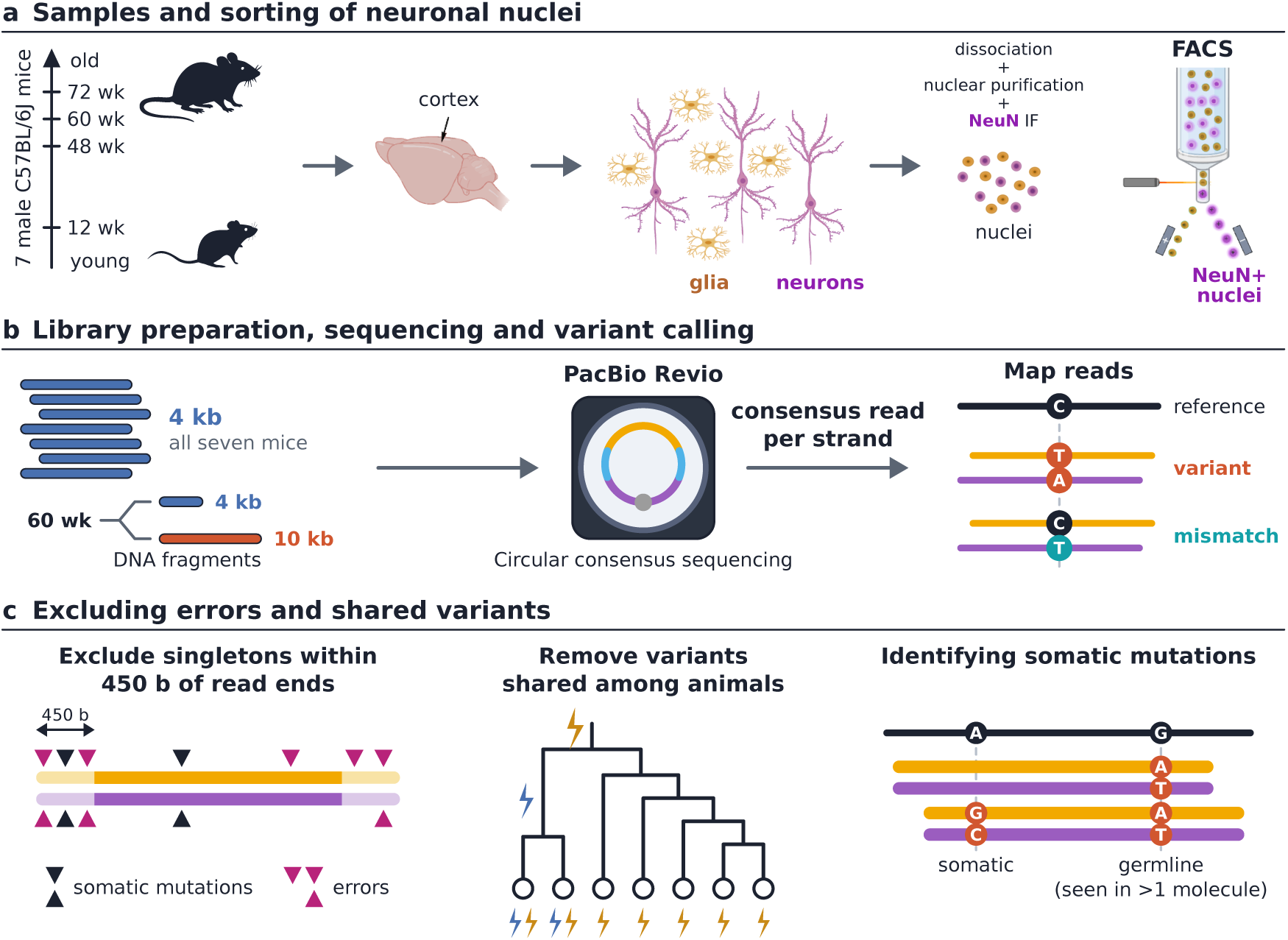
Overview of the experiment and analysis pipeline. a, Samples and sorting of neuronal nuclei. Neuronal nuclei were sorted from the cortex of seven male C57BL/6J mice aged 12, 48, 60 or 72 weeks. The cortex-to-FACS images were made with BioRender. b, Library preparation, sequencing and variant calling. A library of 4 kb fragments (blue) was prepared from each animal, and a second library of 10 kb fragments (red) from the 60-week-old animal. A separate consensus read was built for each strand of every molecule (orange and purple). The strand consensus reads were aligned to the reference genome. A variant is called at a site at which the two strand consensus reads of a molecule carry complementary bases that differ from the reference base; a mismatch is called at a site at which only one of the two reads differs from the reference base. c, Excluding errors and shared variants. Left, singletons in the 450 bases of each read end (pale segments) are excluded; errors (magenta triangles) are enriched there relative to mutations (black triangles). Middle, variants shared among animals are removed; in the schematic genealogy, circles are the seven animals and lightning bolts are mutations inherited by two (blue) or all (yellow) of them. Right, variants carried by two or more molecules are classified as germline, and singletons as somatic. More details about the filters are provided in Supplementary Fig. 1.

Our goal was to maximize the coverage of the genome, while keeping error rates low enough relative to *de novo* mutation rates per cell. To this end, we sought to increase the number of subreads used to build a consensus sequence, by aiming for shorter DNA fragments. In this regard, there is a trade-off: the advantage of shorter fragments is that they are read more times, and thus each consensus sequence is built from more subreads. However, each molecule includes fewer bases, and therefore aiming for shorter fragments leads to lower coverage. To explore this trade-off, we aimed for 4–5 kb fragments for most libraries and 10 kb for one library, compared with the target fragment length of 15–20 kb in the default whole-genome sequencing protocol^30,31^, and median 1.7 kb in the HiDEF approach^29^. In practice, we obtained reads with mean lengths of 3.6 to 4 kb for seven libraries and 10.9 kb for the longer read library (Supplementary Fig. 2), which corresponded to an average of 17–21 and 7 passes per strand, respectively (see Methods).

Another change relative to the default PacBio HiFi protocol is that we built a separate consensus for the two strands of each molecule^29^. We conditioned on each strand consensus being supported by 5 or more subreads (which was true of 98% of the reads in the short libraries and 91% of the reads in the long library). A single nucleotide variant was then called if both strands carried the same non-reference allele. In turn, mismatches between the two strands were used to model sequencing and library preparation errors, informing our approach to mutation calling (see Methods).

#### Estimating error rates

Of interest to us are heterozygous sites in neurons. That set of variants includes inherited germline mutations that arose since the common ancestor with the reference genome sample; *de novo* germline mutations; somatic mutations that arose in development (e.g., in dividing progenitors before neurons exit the cell cycle to become postmitotic); and finally mutations that arose in the postmitotic neuron. A heterozygous germline variant, whether inherited or *de novo*, is carried by a given molecule with probability 1*/*2. At a site covered by *n* molecules, it is therefore carried by exactly one molecule with probability *n/*2*^n^*, which is at most 1% given our minimum coverage of 10 molecules (see Methods). By contrast, most somatic mutations are carried by few molecules^32^. Notably, mutations arising in postmitotic neurons are carried by at most one molecule. Variants carried by a single molecule (henceforth “singletons”) are therefore central to our analysis. They are also the most prone to false discoveries, owing to errors that arise in the library preparation or sequencing. For this reason, we first applied a series of quality checks and filters.

First, because of end repair and A-tailing, we expected an excess of library preparation errors at the end of sequencing reads^28^. The density of singletons is indeed enriched near the ends of molecule consensus reads, within *∼*450 base pairs (bps) of either end, whereas variants carried by two or more molecules are not (Supplementary Fig. 3). We therefore discarded singletons within 450 bases of either end of the molecule, but kept variants carried by two or more molecules irrespective of their position in the read (see Methods).

To estimate how much library preparation and sequencing errors contribute to singletons in the remaining interior portion of the reads, we analyzed mismatches between the two strands of a molecule. First, we considered the mismatch rate as a function of the number of passes. We found that it decreases rapidly: reads supported by 5 passes have a mismatch rate of 3.7 *×* 10*^−^*^5^ per base, which then plateaus at around 9 *×* 10*^−^*^7^ at high (*>*35) numbers of passes (Supplementary Fig. 5). We interpret the apparent asymptote as the upper bound on the rate of single-strand library preparation errors (Supplementary Fig. 5); it is an upper bound because a subset of the mismatches supported by a large number (e.g., *>*35 passes) might also reflect DNA damage that accumulated *in vivo*. Given that only 15% of the overall mismatch rate of 5.9 *×* 10*^−^*^6^ per base persists at high number of passes, most errors appear to be due to misincorporation of nucleotides by the sequencing polymerase or errors in base calling.

We then looked at the mismatch spectrum, classified into 192 single base substitution types based on the trinucleotide context (Supplementary Fig. 6; Methods). This analysis revealed that the most frequent substitution types were rarely matched by their reverse complement on the opposite strand. Assuming errors on the two strands are independent, we estimated the rate at which both strands carry the same error to be 6 *×* 10*^−^*^11^ per bp (see Methods). We also used the fraction of subreads that agree with the consensus in order to independently estimate the sequencing error rate, which yielded an estimate of 2.7% per base per subread (see Methods). In sum, variant calls based on two independent strand consensus reads, each with at least 5 passes, should have few errors: over the callable bps of the eight libraries, we expect 10 false discoveries among the 2,289 singletons (0.4%).

#### Identifying somatic mutations

Next, we sought to distinguish mutations that arose in postmitotic neurons from those that arose earlier in development or were inherited. As a first step, we removed all mutations found in more than one animal, regardless of the number of molecules carrying the non-reference allele (see Methods). Among remaining mutations, those carried by one or two molecules were in excess of the binomial expectation for germline variants (Supplementary Fig. 7). Using a mixture model, we estimated that 99.9% of singletons are somatic mutations (2.2 germline mutations expected among the 2,289 singletons). By contrast, only 14% (95% CI: 12% to 16%) of the variants carried by two or more molecules were inferred to be somatic, mostly those carried by two (of which 89% were estimated to be somatic) or three (52%) molecules (Supplementary Fig. 7). On that basis, we classified singletons as somatic and mutations carried by two or more molecules as germline; the latter set may also include mutations that arose very early in development, which are carried by a large fraction of cells. However, given that each site is covered by only 10–49 molecules sampled from *∼*10^6^ nuclei per animal, most mutations that arose in development are expected to be found in a single molecule^33,34^.

This approach allows us to reliably call somatic mutations without a matched tissue sample, unlike standard methods ^35^. Given our coverage, we estimated that labeling all singletons as somatic mutations misclassifies 0.19% of the heterozygous germline variants (corresponding to 0.10% of singletons). In an outbred species, with orders of magnitude more heterozygous sites, these incorrect assignments would swamp the true somatic mutation signal. However, a related approach should be feasible, as shown by the studies of non-crossover recombination events in primates^26,27^. In species with heterozygosity *h*, a read of length *l* spans *hl* heterozygous sites on average, so most long reads can be assigned to one of the two parental haplotypes. A germline variant will be present on every read of one haplotype, whereas a somatic variant will only be present on a subset. Within each set of phased reads, at any site covered by at least two reads from the same haplotype, somatic mutations can then be called without relying on the number of molecules carrying the variant. Given sufficient coverage, phasing would therefore make our approach applicable to outbred species, without a need for matched tissue samples.

To further test the reliability of the mutation calls, we compared the two libraries obtained from the 60-week-old mouse, with average read lengths of 3.6 versus 10.9 kb. Of the non-reference alleles carried by two or more molecules, 97% (365 of 378) were shared between the two libraries (Supplementary Fig. 8). By contrast, and as expected, singletons in the 60-week-old mouse were almost never found in the two libraries (1 of 441), but their estimated somatic mutation rates were in agreement (Supplementary Fig. 8). The high concordance provides further evidence that our calls are reliable. The comparison also shows that the tradeoff between error rate and coverage is better served by our longer library of 10.9 kb reads. In what follows, we analyze the two libraries for the 60-week-old mouse jointly.

### Effect of age on the number of point mutations

Coverage varies along the genome and between libraries (Supplementary Figs. 9 and 10). To estimate the somatic mutation rate, we therefore divided the number of somatic point mutations by the number of opportunities we had to detect them, i.e., the number of molecules covering each site that passed all filters, summed over all sites (see Methods). Regressing the mutation rate on age, we found a significant increase of 25.9 point mutations per diploid genome per year (by ordinary least squares regression, 95% CI [17.7–34.1], *P* = 4.6 *×* 10*^−^*^4^; Fig. 2). As expected, if the minimum coverage is decreased from 10 molecules to 5, the number of germline and early developmental mutations among the singletons increases and the intercept rises to 62 from 49 point mutations per diploid genome. However, it has little effect on the slope (for which the estimate becomes 27.9, 95% CI [18.9–36.9], instead of 25.9; Supplementary Fig. 11).

**Figure 2.**
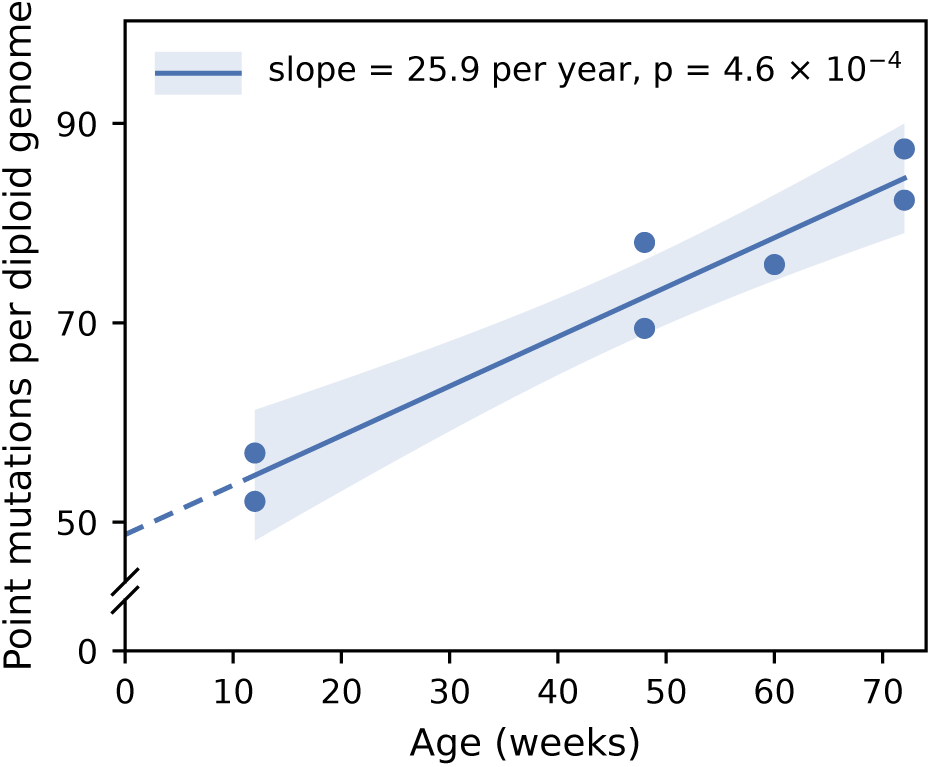
Somatic point mutations in mouse neurons as a function of age. Shown is the number of somatic point mutations per callable bp multiplied by the diploid genome size (5.28 Gb). Blue dots are estimates for the seven animals; the two libraries of the 60-week-old mouse are pooled. The line is the ordinary least squares regression fitted to the blue markers (slope: 25.9 SNVs per diploid genome per year; 95% CI: 17.7 to 34.1; *P* = 4.6*×*10*^−^*^4^, two-sided *t* test; *R*^2^ = 0.93). The shaded area is the 95% confidence interval for the regression line.

Previous studies of mouse neurons were not well powered to detect an age effect, owing to a lack of variation in ages ^19,21^. Nonetheless, in the study of mouse cerebellar neurons, which is comparable in design to ours, their point estimate of the mutation rate is highly consistent with our prediction for that age (19 months; Supplementary Fig. 12). Moreover, the types of mutations identified in our study are similar to theirs^19^, albeit statistically significantly different (the cosine similarity to a 96-channel mutation spectrum is 0.94; *P* = 10*^−^*^4^; Fig. 3); such subtle differences in spectra between neuronal cell types are also seen in humans ^36^. The mutations we identified are relatively well modeled by mouse signatures mSBS1 and mSBS40^37^ as well as by human signatures found in human neurons^15,17^ (cosine similarity 0.94 and 0.93, respectively; Fig. 3b). Also as reported for human neurons^17^, mutations in mouse neurons are enriched in accessible chromatin regions (1.4-fold; one-sided *P* = 0.011; Fig. 3c). Thus, the neuronal mutations that we identified match properties of those obtained using other, bespoke approaches.

**Figure 3.**
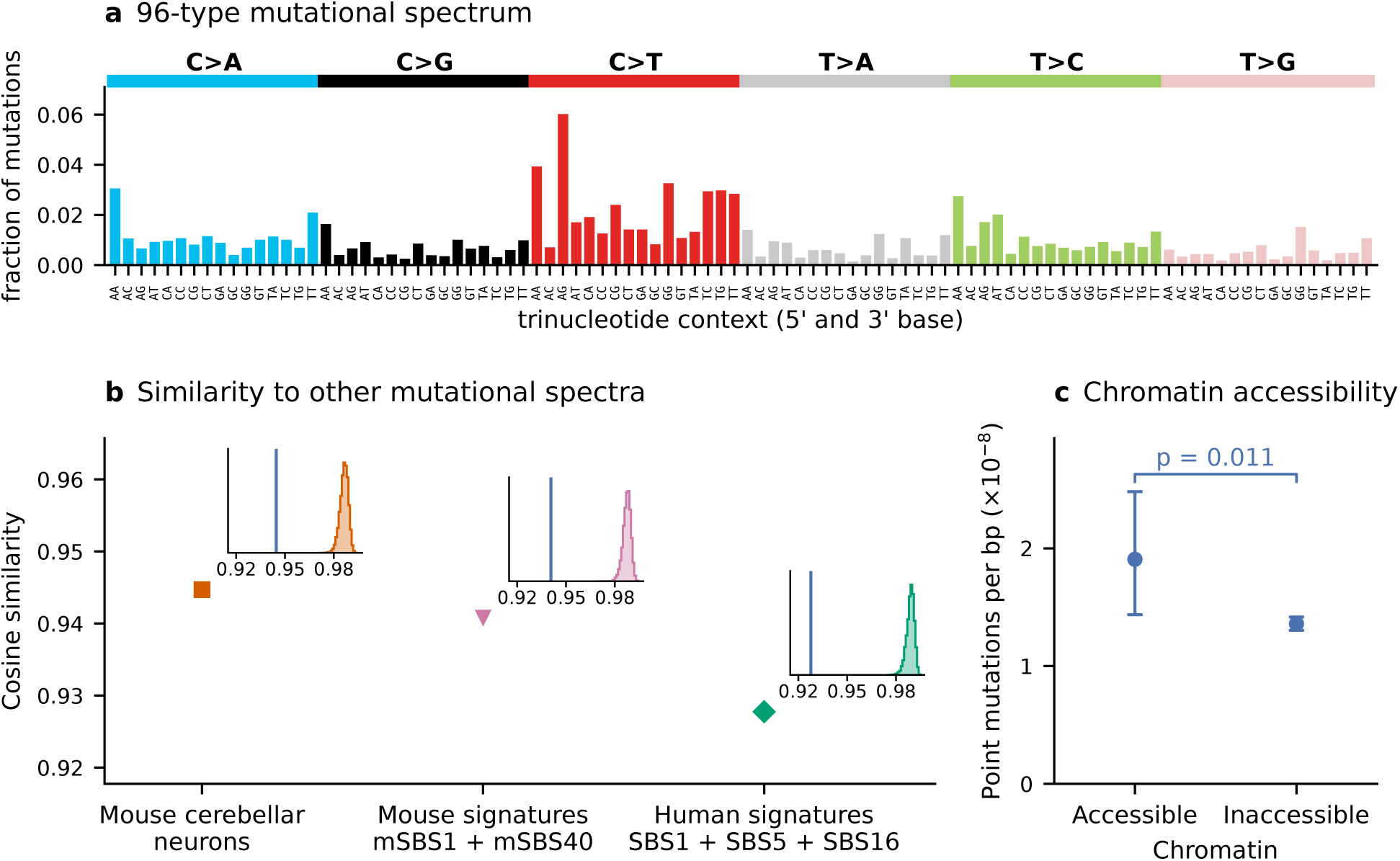
Spectrum and genomic distribution of point mutations in mouse cortical neurons. a, 96-type mutational spectrum of the 2,289 point mutations, pooled over the seven animals after correcting each animal for the trinucleotide composition of its callable bps relative to the T2T autosomes. b, Cosine similarity of our spectrum to that of cerebellar neurons of mice fed *ad libitum* ^19^ and to the best-fitting combinations of two mouse signatures (17% mSBS1 + 83% mSBS40) and of three signatures seen in human neurons (13% SBS1 + 87% SBS5 + 0% SBS16; COSMIC v3.4) (see Methods). Note that the y-axis starts at 0.92. Insets show the cosine similarity of our neuronal mutations (blue line) versus the distributions of cosine similarities expected by chance, given sampling error (see Methods). c, Point mutations per callable bp in accessible and inaccessible chromatin. Error bars are exact 95% Poisson confidence intervals. The *P* value is from a one-sided, exact conditional test for two Poisson rates (see Methods).

### Implications

Human cortical neurons were found to accumulate 16 to 23 point mutations per diploid genome per year ^14–17^, estimates that overlap with our 95% CI (Fig. 2). Thus, our estimated rate in mouse neurons is only 1.1–1.6-fold that reported in humans. This observation stands in contrast to the 17-fold difference in colonic crypts^9^ and 13-fold difference in germline mutation rates^38^. Neurons therefore do not follow the scaling of yearly mutation rates with lifespan seen in the colon or with generation time seen in the germline. In this regard, we note that during the preparation of our manuscript, a preprint was posted that used single-cell, whole-genome amplification to call point mutations in cortical neurons and found a similar age effect to humans in eight BALB/c *×* C57BL/6 F1 hybrid mice, as well as chimpanzees, rhesus macaques, marmosets and ferrets^39^. In contrast to our findings for mice, however, the spectrum of mutations identified in non-human primates was markedly different from that in humans (and no comparison was reported for mice) ^39^.

Whether the difference between germline and intestinal crypts on the one hand and neurons on the other reflects more general rules about cell types or not remains to be explored. A fruitful approach to this question may come from application of PacBio HiFi sequencing to call point mutations at single-cell resolution. As we show, the approach is sensitive enough to detect a subtle effect of age and reliable enough to recover properties of mutations identified by alternative approaches. The use of an off-the-shelf technology to characterize mutations at single-cell resolution opens the door to the study of mutations across cell types and across the tree of life and should therefore enable longstanding questions about mutagenesis and the selection pressures that shape mutation rates to be addressed ^5,6,40,41^.

## Acknowledgments

We thank Maya Fridrikh and the Genomics Core Facility at the Icahn School of Medicine at Mount Sinai for assistance with sequencing and with the related Methods section, as well as Sara Bizzotto, Gilad Evrony, Marc de Manuel, Mikkel Schierup and Guy Sella for helpful discussions. This work was funded by a Research Initiative in Science and Engineering grant to MP and FP from Columbia University.

## Methods

### Mouse samples

Male C57BL/6J mice (JAX #000664) were analysed at 12, 48, 60 and 72 weeks of age (*n* = 2 per age group). One 60-week-old mouse was later excluded, and two libraries were prepared from the remaining 60-week-old mouse (see Library preparation). Mice were housed under a 12h light/dark cycle with *ad libitum* access to food and water. All procedures were approved by the Columbia University IACUC committee.

### Isolation of neuronal nuclei from mouse cortex

Neuronal nuclei were isolated from mouse cortex using a protocol adapted from Liu et al.^29^ . Cortices were dissected in ice-cold PBS, cut into *∼*5 mm^3^ fragments, transferred to cryovials, snap-frozen in liquid nitrogen, and kept at *−*80 °C until processing. All buffers and a large Dounce homogenizer (Sigma-Aldrich, D9938) were chilled on ice before use. Frozen tissue from each animal was placed in 9 ml of cold lysis buffer (0.32 M sucrose, 10 mM Tris-HCl pH 8.0, 5 mM CaCl_2_, 3 mM magnesium acetate, 0.1 mM EDTA, 1 mM DTT, 0.1% Triton X-100) and homogenized on ice with 20 strokes of pestle A followed by 20 strokes of pestle B. The lysate was carefully overlaid onto 7.4 ml of sucrose cushion buffer (1.8 M sucrose, 10 mM Tris-HCl pH 8.0, 3 mM magnesium acetate, 1 mM DTT) in an ultracentrifuge tube and spun at 10,000 rpm for 1 h at 4 °C. After the supernatant was discarded, 500 µl of resuspension buffer (1*×* PBS containing 3 mM MgCl_2_) was added over the nuclear pellet. The pellet was left on ice for 10 minutes and then gently resuspended.

For staining, 1.2 µg of NeuN-Alexa Fluor 647 antibody (Abcam, ab190565) and 0.7 µl of DAPI (2 mg/ml stock) were diluted in 400 µl of staining buffer (3% BSA in resuspension buffer) and mixed by gentle inversion. The staining mixture was added to the nuclei suspension at a ratio of 400 µl per 1 ml of suspension (final DAPI concentration *∼*1 µg/ml, 1:2000), and the sample was transferred to a 2 ml DNA LoBind tube (Eppendorf, 022431048) and incubated with rotation for 30 minutes at 4 °C. NeuN^+^ nuclei were isolated by fluorescence-activated nuclei sorting on a Sony LE-MA900 sorter, gating sequentially on DAPI^+^ singlets and then NeuN^+^ events. Sorted nuclei were collected into 2 ml DNA LoBind tubes containing 30 µl of resuspension buffer, and tubes from the same sample were pooled into a 15 ml DNA LoBind tube (Eppendorf, 0030122208). To stabilize the nuclei during pelleting, an equal volume of 80% glycerol was added, giving a final concentration of 40%. Nuclei were then pelleted at 500 *× g* for 10 minutes at 4 °C, the supernatant was removed, and the pellets were immediately frozen at *−*80 °C.

### Library preparation

Library preparation and sequencing were performed at the Genomics Core Facility at the Icahn School of Medicine at Mount Sinai. High molecular weight (HMW) gDNA was extracted from 8 samples of 1.5 million neuronal nuclei in *∼*100 µL of 40% glycerol using the Qiagen MagAttract HMW DNA kit (67563), per manufacturer protocol. Briefly, Proteinase K, RNase A, and lysis buffer were added to the sample, which was then incubated at room temperature. MagAttract beads were added along with a beads buffer and the samples were incubated with the beads at 1400 rpm for 3 minutes. Beads were washed twice with 2X volume of a Qiagen wash buffer, then twice with an ethanol wash buffer, and finally twice with water. 100–200 µL of elution buffer was added to each sample for a total of 520–3000 ng recovery with average DNA lengths between 40 kb and 70 kb.

Owing to low DNA yield, one 60-week sample was excluded, leaving seven samples. From each, one library targeting 5 kb fragments (“short”) was prepared; from one 60-week sample, an additional library targeting 10 kb fragments (“long”) was also prepared, for a total of 8 libraries from 7 mice. For 7 short libraries, 180–520 ng of HMW gDNA obtained via MagAttract protocol were sheared to around 5 kb using a modified Covaris G-tube shearing protocol (pn_010154). Namely, gDNA was sheared at the highest recommended centrifuge speed given the input mass, diluted, then further sheared at this speed. For samples that returned longer than desired, dilution and re-shearing were repeated until 5 kb length was reached. The long library with 1.3 µg of HMW gDNA was processed in parallel using standard shearing procedures to achieve a length of 10 kb. Sheared DNA was cleaned using 1X SMRTbell cleanup beads with roughly 80–90% recovery and assessed for quantity using the Qubit 1X dsDNA HS assay and length using the Agilent BioAnalyzer.

All libraries were prepared for PacBio single-molecule real-time (SMRT) sequencing mostly as per manufacturer’s instructions using the “Preparing whole genome and metagenome libraries using SMRTbell prep kit 3.0” protocol (PacBio 102-166-600), with exceptions for the shorter DNA length and lower input mass. For 5 kb (10 kb) samples, 140–460 ng (1.2 µg) of sheared, cleaned DNA was used as input. Single-stranded DNA ends were removed and nicks in the DNA were treated enzymatically. DNA blunt ends were repaired, and the template ends were A-tailed (PacBio 102-182-700). SMRTbell barcoded overhang adapters were ligated to each sample (PacBio 102-009-200), after which unligated or damaged templates and unligated adapters were digested in a nuclease treatment step. The final libraries were cleaned using a 1.0X SMRTbell beads cleanup step for final average lengths between 4.6 and 5.6 kb. The longer library was additionally size-selected on Sage Science BluePippin from 7 to 15 kb (High Recovery), for a final average size of 11.7 kb. The final, size-selected libraries were quantified using the Qubit 1X dsDNA HS assay and checked for length using the BioAnalyzer.

### DNA sequencing and read processing

Libraries were separately annealed to PacBio Revio sequencing primer and bound to the Revio SPRQ polymerase for sequencing on individual 25M SMRT cells on the Revio system, in which each molecule is sequenced in its own zero-mode waveguide. The Revio performs circular consensus sequencing (CCS) on-instrument (ccs v8.2.0; Instrument Control Software v13.3) and retains only CCS reads with high accuracy (*>*Q20), at least 3 polymerase passes, and acceptable barcode quality scores as HiFi reads. Runs were initiated using custom .csv file upload to achieve CCS by strand, in which a separate consensus is built for each strand of a molecule so that each zero-mode waveguide yields two strand consensus reads, and to output tags such as the number of polymerase passes for each strand’s consensus read (np), the number of subread alignments covering each base in the consensus read (sa), the number of subreads matching that base (sm), and the number of subreads mismatching it (sx). We note that PacBio Revio does not currently provide methylation information for by-strand consensus sequences. For a full list of output tags, see Supplementary Table 1. The number of reads and molecules, read lengths and numbers of passes for each library are given in Supplementary Table 2.

### Alignment

Reads were aligned to the telomere-to-telomere (T2T) assembly of C57BL/6J (C57BL_6J_T2T_v1, GenBank GCA_964188535.1) ^42^. This assembly contains the 19 autosomes and 218 unplaced scaffolds but not the sex chromosomes or the mitochondrial genome. We added chromosomes X and Y and the mitochondrial genome from GRCm39. The additional chromosomes and the unplaced scaffolds served only as decoys, to capture reads from sequences missing from the autosomal assembly that might otherwise align to the autosomes. All downstream analyses were restricted to the 19 T2T autosomes.

The forward and reverse strand consensus reads were aligned independently with pbmm2 (v26.1.99; Pacific Biosciences), a wrapper around minimap2^43^, using the HiFi preset (--preset CCS) and an index built with the same preset. Alignments were sorted by pbmm2 (--sort) and indexed with samtools (v1.22.1) ^44^.

### Variant calling and filtering

We called somatic variants by comparing the two strand consensus reads of each molecule with each other and with the reference. A point mutation was called at a site if the bases of both strand consensus reads of a molecule passed the filters below and were complementary to each other but differed from the reference. Filters were applied to strand consensus reads, to molecules and to bases before calling, and to sites and variants afterwards (Supplementary Fig. 1).

#### Read filters

We kept only primary alignments to the T2T autosomes with mapping quality of 60 (the maximum assigned by pbmm2). A strand consensus read was also discarded if it was built from fewer than 5 passes (Supplementary Fig. 13), if its length fell outside the mean *±* 2 s.d. of the read-length distribution of its library (Supplementary Fig. 2), or if its divergence from the reference exceeded 1%. Divergence was defined as 1 *− M/*(*M* + *X* + *I* + *D*), where *M* , *X*, *I* and *D* are the numbers of matched, mismatched, inserted and deleted bases in the alignment.

#### Molecule filters

The two strand consensus reads of each molecule were paired by the zero-mode waveguide number in their read names. A molecule was kept only if both of its strand consensus reads passed the read filters and aligned to the same genomic location. We then removed molecules for which the two strand consensus reads differed in length by more than 62–70 bases in the short libraries and 125 bases in the long library (the library’s 99th %-tile). We also removed chimeric molecules, in which part of either strand consensus read aligned to a second location in the genome (a supplementary alignment). We further removed molecules whose two alignments differed most in the reference genome interval they covered, i.e., when the number of sites covered by only one of the two alignments exceeded the library’s 99th %-tile. Library-specific thresholds and the number of reads and molecules remaining after each filter are given in Supplementary Table 3.

#### Base filters

Within each strand consensus read, a base was used only if three conditions held: its Phred base quality was at least 40; at least five subreads matched it (sm *≥* 5), and these made up more than 80% of the subreads aligned at that base (sm/(sm + sx) *>* 0.8); and it lay more than three bases from an insertion or deletion in the alignment of that read. Sites at which the reference base or either of its neighbours was N were excluded.

#### Masked sites

We masked three kinds of sites: those in simple repeats, low-complexity sequence and satellite DNA (RepeatMasker classes Simple_repeat, Low_complexity and Satellite, as annotated for this assembly in the UCSC Genome Browser; RepeatMasker v4.1.4, Dfam 3.6). Only three sites had more than one alternative base in a library, all of them in the masked repeats. The mask covers 296.1 Mb, or 11.2% of the autosomes.

#### Coverage filter

The coverage of a site is the number of molecules for which the two strand consensus reads passed the read, molecule and base filters and carried complementary bases. We used the coverage in the untrimmed reads. The mean coverage of autosomal sites covered by at least one molecule was 11.4–14.1 molecules across libraries (Supplementary Fig. 9). We kept sites with coverage of at least 10 molecules to be able to reliably distinguish somatic (expected at low frequencies) and constitutive variants (expected at frequency 50%). Sites with unusually high coverage may reflect collapsed repeats or alignment errors; we therefore excluded sites with coverage above the 99th %-tile of each library (21–25 molecules in the short libraries and 28 in the long library). After masking and filtering, we are left with 58.9–74.0% of the autosomes for each sample (Supplementary Table 4).

### Errors at the edges of consensus reads

Variants called near the ends of strand consensus reads are enriched for errors, which likely arise during end repair and A-tailing ^15,29^. To infer the distance over which this enrichment occurs, we compared the density of singletons (non-reference alleles carried by a single molecule in a single mouse), for which we expect a mixture of mutations and errors, with the density of a control set among which errors should be negligible, non-reference alleles carried by at least two molecules in a library. We pooled all libraries to increase power.

In reads of at least 3 kb, we counted the singletons, control variants and callable bps at each distance *d* = 6*, . . . ,* 1,500 bases from the nearest read end (the first five bases were excluded because there, the aligner clips mismatched bases). We defined the interior of a read as sites more than 1,500 bases from the end of a read of at least 4 kb. For each library *l*, we estimated the density *r*_0_*_,l_* in the interior as the number of singletons divided by the number of callable bps. The number of singletons expected at distance *d* in the absence of edge errors is then 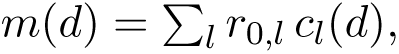, where *c_l_*(*d*) is the number of callable bps of library *l* at distance *d*. We found that, as expected, the density of control variants does not depend on the distance from the read end (*χ*^2^ test across ten 150-base bins, *P* = 0.39; Supplementary Fig. 3b), whereas singletons are enriched near the read ends (same test, *P* = 4 *×* 10*^−^*^96^; Supplementary Fig. 3a).

To estimate how many edge errors remain after trimming *t* bases from each read end, we modeled the number of singletons at distance *d* as *s*(*d*) = *m*(*d*) + *a*(*d*), where *a*(*d*) is the number of edge errors at distance *d*, with no assumed form for the dependence on *d*. The only estimated quantities are therefore the interior densities *r*_0_*_,l_*. The estimated number of edge errors left after trimming is

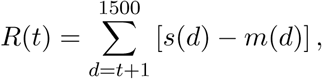

and the false discovery rate (FDR) is *R*(*t*)*/K*(*t*), where *K*(*t*) is the number of singletons left in the same reads after the trim, more than *t* bases from both ends. 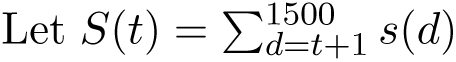 be the number of singletons from *t* + 1 to 1,500 bases from the nearest read end. The fraction of the singletons within 1,500 bases of a read end that are edge errors is then *f* = *R*(0)*/S*(0). Supplementary Fig. 4 plots the cumulative fraction of singletons against the distance from the read end, together with the expectation if all edge errors were to lie at the read end, *f* + (1 *− f* )*M* (*d*), where *M* (*d*) is the cumulative fraction of *m*(*d*). At any distance *d*, the gap between the two curves is *R*(*d*)*/S*(0).

We trimmed *t* = 450 bases from each end of the strand consensus reads, as the two curves converge beyond that distance (Supplementary Fig. 4). We note that a non-reference allele can be carried by a single molecule in a trimmed read and by other molecules within the 450 bases removed from their ends. We therefore counted a non-reference allele as a singleton only if it remains a singleton when considering the untrimmed reads of other molecules. We removed molecules carrying more than 2 singletons from further analyses.

After the trimming, edge errors account for a small fraction of the singletons: the estimated FDR is *R*(450)*/K*(450) = 0.6%, with a one-sided 95% upper confidence bound of 5.4% (using a normal approximation, assuming that the numbers of singletons 451–1,500 bases from the read ends, and in the read interiors, are Poisson-distributed).

### Somatic and germline mutations

We classified singletons as somatic mutations (2,289 in total, 211–605 per animal) and variants carried by two or more molecules of an animal (pooling the two libraries of the 60-week-old mouse) as germline or early embryonic mutations (1,350 in total, 147–264 per animal). Neither set had unusually high or low coverage: the median untrimmed coverage was 13–16 molecules at the somatic SNVs, 13–15 at the germline candidates and 13–16 over all unmasked sites with an untrimmed coverage of at least 10 (Supplementary Fig. 10).

To assess the reliability of this classification rule, we estimated the fraction of somatic variants among those seen in at least two molecules. For each variant, *k* of the *n* molecules covering the site carry the alternative allele. For this test, we used variants with *n ≥* 10 and 2 *≤ k ≤ n −* 1; 1,343 in total. We modeled the distribution of *k* as a mixture. For germline variants, we used a binomial distribution 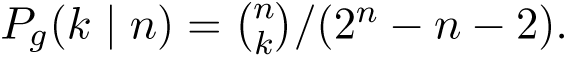. For somatic variants, we used the site frequency spectrum of a pure-birth process, which is proportional to 1*/*[*k*(*k −* 1)] ^33^. For 2 *≤ k ≤ n −* 1, *P_s_*(*k | n*) = (*n −* 1)*/*[(*n −* 2) *k*(*k −* 1)]. We estimated the somatic fraction *p* to be 0.138 (95% CI: 0.116 to 0.163) by maximum likelihood using expectation-maximization ^45^. A germline variant at a site covered by *n* molecules is carried by a single molecule with probability *n/*2*^n^*, and by 2 to *n −* 1 molecules with probability 1 *−* (*n* + 2)*/*2*^n^*. From these probabilities and the fitted mixture, 2.2 germline variants are expected among the singletons, corresponding to 0.10% of singletons and 0.19% of heterozygous germline variants.

### Estimation of the somatic mutation rate

For each animal, the somatic mutation rate per bp is defined as the ratio of the number of somatic mutations (numerator) and the number of callable bps (denominator). The number of callable bps of a library is given by the coverage summed over its callable sites (excluding the read edges, in which we do not call singletons) (Supplementary Table 4). For the 60-week-old mouse, the number of mutations and the number of callable bps were summed over its two libraries (see Concordance between the short and long libraries of the 60-week-old mouse).

### Estimation of the sequencing error rate

We estimated the error rate per bp per molecule from mismatches between the two strand consensus reads of each molecule, using the trimmed reads. A mismatch is a site at which the bases of both strand consensus reads passed all read, molecule and base filters (see Variant calling and filtering), yet one of the reads carried the reference base and the other a non-reference base. We ignored the small possibility that library preparation errors may propagate to the other strand. Moreover, we treated every mismatch as a sequencing error, although some may reflect unrepaired single-strand DNA damage; in that regard, the resulting estimates of error rates are upper bounds. We based our estimates on the unmasked sites without the coverage filter. Across the eight libraries, there were 2,244,072 mismatches over 3.81 *×* 10^11^ bases, counted once for each strand (average of 5.9 *×* 10*^−^*^6^ mismatches per base per strand).

We estimated error rates separately for each of the 192 strand-oriented trinucleotide substitution types and for each class of passes of the divergent read, correcting for the number of opportunities (Supplementary Table 5). Each mismatch was recorded as read 5*^′^ →* 3*^′^* on the strand carrying the non-reference base. We separated the classes of passes because the mismatch rate depends on the number of passes (Supplementary Fig. 5), and the two orientations because mismatches in which a pyrimidine was changed on the read carrying the non-reference base had a different spectrum from those in which a purine was changed (Supplementary Fig. 14).

For a sequencing error to be erroneously called a somatic mutation, both strand consensus reads of a molecule must carry the same non-reference base at the same position. Assuming that errors on the two strands are independent, we estimated the rate of each of the 96 substitution types as the product of the rates of its two reverse-complementary substitution types in the same class of passes. For each library, we averaged these rates over the classes of passes, weighting each class by the mutational opportunities in that class and trinucleotide context. We then multiplied each rate by the callable bps in that context (Supplementary Table 6) and summed over the 96 substitution types. This approach led us to estimate 10.0 false discoveries in total, which would represent 0.44% of the 2,289 somatic point mutations: 0.6–0.8 in each short library (0.2–0.3% of its somatic mutations) and 5.2 in the long library (1.3% of its 410), in which consensus reads are based on fewer passes.

We also estimated the error rate of individual subreads from the number of subreads matching (sm) and mismatching (sx) each base of a strand consensus read (Supplementary Table 1). In each library, we sampled 10 million bases at random from 100,000 strand consensus reads, excluding the first and last 450 bases of each read, and divided the number of subreads that disagreed with the consensus base by the total number of subreads at these bases. This approach led us to estimate an error rate of 2.7% per base per subread over the eight libraries (2.2–3.0% in each library).

### Concordance between the short and long libraries of the 60-week-old mouse

To check that the pipeline gives consistent results for libraries with different distributions of molecule lengths, we compared the short and long libraries of the 60-week-old mouse, which were prepared from the same sample. We used only the sites callable in both libraries, i.e., 1.30 Gb (49.4% of the autosomes).

First, we asked whether non-reference alleles carried by at least two molecules in either library were detected in both libraries (Supplementary Fig. 8a). We included variants in the untrimmed reads, since a variant carried by two or more molecules is unlikely to be an artefact. Of the 378 such variants, 13 are seen in only one library (Supplementary Fig. 8a), all carried by two or three molecules (VAF 0.06–0.12). For one of the 7 variants seen only in the long library, a molecule of the short library carried the variant but had been removed by the read-length filter.

To test whether more non-reference alleles were seen in only one library than expected by chance, we assumed that a molecule has the same chance of carrying a given allele in either library. For a variant carried by *k* of the *n* molecules covering its site in the two libraries, the number of carrying molecules in one of the libraries then follows a hypergeometric distribution, with one modification: since we require *k >* 1, a variant with *k* = 2 will only be included if both molecules come from the same library. Summing over the 378 variants, we expect 9.3 variants to be seen only in the long library and 5.5 only in the short library. The number of variants seen in only one library is a sum of independent outcomes with different probabilities, so follows a Poisson binomial distribution. The probability of at least as many variants as observed was 0.48 for the short library (6 observed) and 0.91 for the long library (7 observed); in other words, the number of variants found in only one library does not differ from what would be expected by chance.

Second, we compared the somatic mutation rates of the two libraries at the same sites (Supplementary Fig. 8b), following the same approach as described in Estimation of the somatic mutation rate. The short library had 161 point mutations over 1.32 *×* 10^10^ callable bps (1.22 *×* 10*^−^*^8^ per bp; 95% CI: 1.04 to 1.42 *×* 10*^−^*^8^), and the long library 279 over 1.96 *×* 10^10^ bps (1.42 *×* 10*^−^*^8^ per bp; 95% CI: 1.26 to 1.60 *×* 10*^−^*^8^). Confidence intervals are exact Poisson intervals. The rate of the long library was 1.17 times that of the short library (95% CI: 0.96 to 1.43; *P* = 0.13, exact conditional test for two Poisson rates). Given that their somatic mutation rates did not differ significantly, we summed their point mutations and callable bps when estimating the rate for that mouse.

### Relationships among the samples

To learn how the seven mice are related, we built a tree from the 501 non-reference alleles carried by at least two molecules, using the 0.63 Gb (24.0% of the autosomes) that were callable in every library, in the untrimmed reads (Supplementary Table 7). Of the 501 alleles, 67 are carried by all seven mice and distinguish the colony from the mouse used for the reference assembly, 69 are carried by two to six mice, and 365 are carried by a single mouse (and thus only contribute to the length of terminal branches). We measured the distance between two animals as the number of allelic differences between them over the 501 sites, with the two libraries of the 60-week-old mouse pooled. We built a neighbour-joining tree ^46^ (Supplementary Fig. 15) with Biopython (v1.88) ^47^ and used the reference genome as the outgroup. Support for each branch was obtained from 10,000 bootstrap replicates over sites^48^.

### Mutational spectra

We classified each point mutation into one of 96 types, defined by the substitution type, written as the change of the pyrimidine of the base pair (C*>*A, C*>*G, C*>*T, T*>*A, T*>*C or T*>*G), and the bases immediately 5*^′^* and 3*^′^*. We corrected counts for each mouse by the trinucleotide opportunities sampled. The corrected counts were then summed over the seven animals.

We compared our spectrum with three other spectra, using the cosine similarity ^49^. Each of these spectra reflects in part the trinucleotide composition of the genome from which it was obtained, so for each comparison we adjusted our counts for the trinucleotide opportunities in that genome. The first is the spectrum of cerebellar neurons from the 12 mice fed *ad libitum* in Grońska-Pęski et al.^19^ . We took point mutations on the autosomes (from their Supplementary Table 5), counting the few mutations that are listed more than once for the same mouse (23 of 2,042) once, as they did (their Supplementary Table 7). To correct for the trinucleotide frequencies, we used the ratio they report for each mouse separately (their Supplementary Table 7). In turn, when comparing our mutations to signatures that Riva et al.^37^ extracted from tumour genomes of mice aligned to GRCm38, or to the COSMIC v3.4 single-base substitution signatures for GRCh37^50^, we used GRCm38 and GRCh37, respectively. For the mouse signatures, we fitted mSBS1 with mSBS5 and mSBS1 with mSBS40, and kept the pair with the higher cosine similarity: 17% mSBS1 and 83% mSBS40. For the human signatures, we fitted the signatures identified in neurons (SBS1, SBS5 and SBS16)^15,17^; in the best fit model, 13% of the mutations were attributed to SBS1, 87% to SBS5 and 0% to SBS16.

To test whether the cosine similarity to our mutation spectrum is lower than expected from sampling error alone, we asked what it would look like if our mutations were drawn from the spectrum to which we were comparing. Specifically, in each of 10,000 simulations, for each of our mice, we randomly drew as many point mutations as observed, adjusted for trinucleotide opportunities of the mouse, with replacement. We then treated the simulated set like the real data and examined the distribution of the cosine similarity over the 10,000 runs. The *P* value is the number of simulations with a cosine similarity as low as or lower than ours, plus one, divided by 10,001.

### Chromatin accessibility

We defined accessible chromatin in cortical neurons from the single-nucleus atlas of the mouse brain of Sziraki et al.^51^ (GEO: GSE212606), which profiled C57BL/6 mice. We pooled the 50,858 nuclei of wild-type mice in the clusters “Cortical projection neurons 1–3” and “Interneurons 1–2”.

The definition of accessible chromatin follows that of Grońska-Pęski et al.^19^ . Specifically, we considered the accessible chromatin to consist in the snATAC-seq peaks of that study in which at least 2% of the pooled nuclei had at least one read (30,850 of 339,951 peaks) and all other sequence in GRCm39 to be inaccessible. We mapped each site of the T2T autosomes to GRCm39 using the alignment chain file from the UCSC GenArk hub for this assembly^52^; sites with no counterpart in GRCm39 (8.5% of the autosomes) were left out. By this definition, accessible chromatin covers 46.0 Mb, or 1.7% of the autosomes. For each region, the mutation rate is the number of point mutations at its sites divided by its callable bps. We tested for a higher rate in accessible chromatin, as has been reported for human neurons ^17^.

### Data and code availability

Mutation calls are provided in Supplementary Table 7. Sequencing reads will be deposited in a public repository and made publicly available prior to publication. The code to reproduce our analyses is available at github.com/matin-saeidi/ccs_somatic_mutations.

## Supplementary Information

### Supplementary Tables

Supplementary Tables 1–7 are provided as a separate spreadsheet.

**Supplementary Table 1**. PacBio tags for strand consensus reads. For each tag, we indicate whether it takes one value per read or one value per base, and how we used it, if at all.

**Supplementary Table 2**. Sequencing output for each library. HiFi reads are the strand consensus reads returned by the instrument, i.e., the sum of the forward and reverse strand consensus reads. Molecules are zero-mode waveguides that yield a consensus read for both strands. HiFi yield is the total number of bases in the HiFi reads. Read lengths (in bases) and numbers of passes are provided per strand consensus read.

**Supplementary Table 3**. Read and molecule filters for each library. The first three rows are statistics for each library: the mean *±* 2 s.d. of read lengths; the thresholds on the difference in lengths between the two strand consensus reads of a molecule (99th %-tile); and the threshold on the number of sites covered by only one of the two alignments of strand consensus reads (99th %-tile). The fourth row and below are the number of strand consensus reads, and then of molecules, that remain after each filter, applied in the order listed, as a percentage of the totals in Supplementary Table 2. The number of molecules used to call variants are reported in the last row.

**Supplementary Table 4**. Callable sites and point mutations for each library. We report the percentage of autosomal sites covered by at least one molecule and the mean untrimmed coverage of these sites. Callable sites are unmasked sites with an untrimmed coverage between 10 molecules and the 99th %-tile in the library (as in Supplementary Fig. 10). The number of callable bps is the denominator used to estimate the somatic mutation rate (see Methods). We also report the number of molecules that were excluded because they carried more than two singletons. The number of somatic mutations are reported in the second to last row. The last row reports the number of non-reference alleles carried by two or more molecules in a single mouse, irrespective of their position in the reads, i.e. the set that we consider germline or early embryonic mutations.

**Supplementary Table 5**. Mismatch rates between strand consensus reads, by substitution type and number of passes. Each row pairs a substitution of a pyrimidine with the reverse-complementary substitution of a purine, both written 5*^′^ →* 3*^′^* as they appear in the read with the non-reference base. For each, we report the number of mismatches, the number of opportunities (i.e., bases in the same trinucleotide context) and their ratio. The last column is the product of the two rates, i.e., the rate at which both strands are expected to carry the same error if errors on the two strands are independent. Rates are given separately by the number of passes supporting the read with the non-reference base; rows labelled “All” pool them.

**Supplementary Table 6**. Callable bps by trinucleotide context. Each row is one of the 32 trinucleotide contexts, i.e., a base and its two neighbours, where a context and its reverse complement are counted as one and written with the pyrimidine in the middle (e.g., ACA includes TGT). For each context, we report the number of callable bps in each library and the number of sites in the T2T autosomes, overall and in unmasked regions, followed by the same quantities as percentages of their column totals.

**Supplementary Table 7**. Somatic mutations, germline or early embryonic mutations and shared non-reference alleles. Each row is a non-reference allele, which we classified as a somatic mutation, a germline or early embryonic mutation, or which is shared among mice. For each variant, we report the chromosome and 1-based position on the T2T autosomes, the reference and alternative bases, the reference trinucleotide, and whether the site was used to build the tree (Supplementary Fig. 15). In columns H–O and P–W, we provide, for each library, the number of molecules carrying the non-reference allele and the coverage at the site, in the untrimmed reads. The number of molecules is left blank when the site was not callable in that library.

## Supplementary Figures

**Supplementary Fig. 1.**
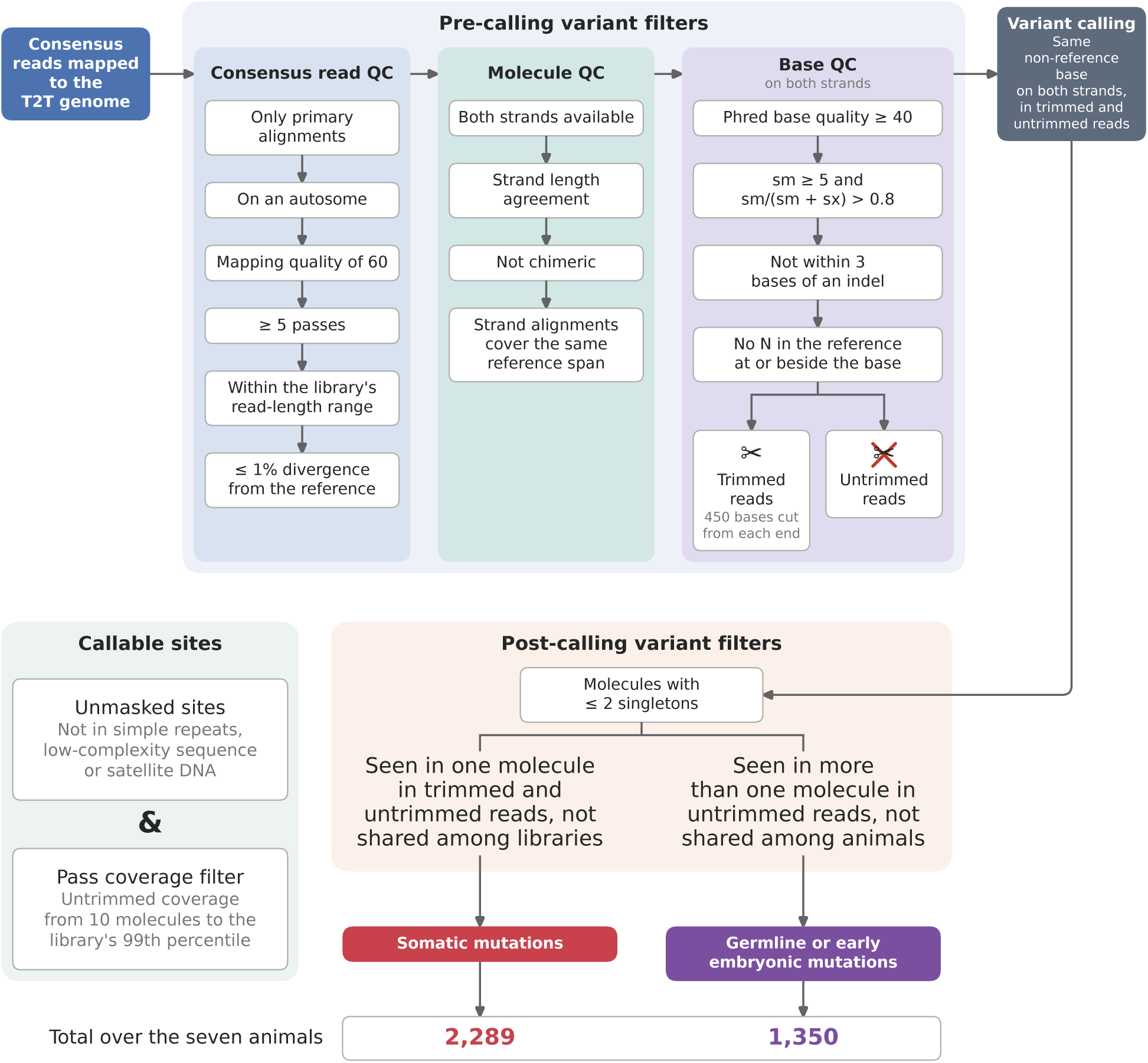
Schematic of the bioinformatic pipeline leading to our variant call sets. Filters are imposed on strand consensus reads (consensus read QC), on molecules (molecule QC) and on the bases of both strand consensus reads (base QC). The last base filter removes 450 bases from each end of every read, yielding “trimmed reads”; variants are called in both the trimmed and the untrimmed reads. After imposing these filters, a variant is called at a site at which the two strand consensus reads of a molecule carry complementary bases that differ from the reference base. Callable sites are the unmasked sites that pass the coverage filter, which uses the coverage in the untrimmed reads. At callable sites, molecules carrying more than two singletons are removed. The remaining variants are classified by the number of molecules carrying them: those carried by one molecule in both the trimmed and the untrimmed reads, and not carried by molecules in other libraries, are considered somatic mutations; those carried by two or more molecules in the untrimmed reads, and called in no other animal, are germline or early embryonic mutations. More details are provided in Methods.

**Supplementary Fig. 2.**
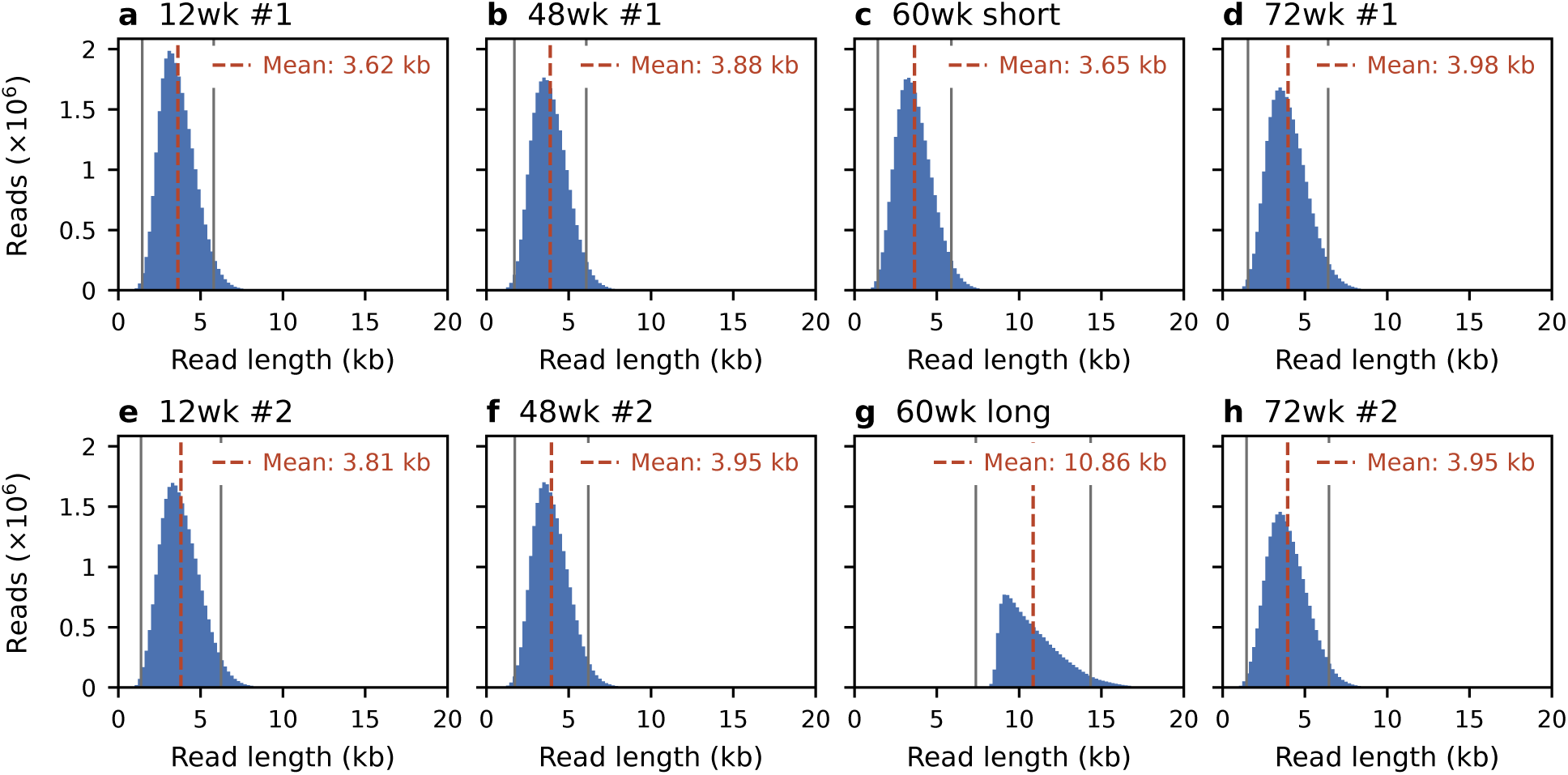
Distribution of read length of HiFi strand consensus reads in each library. Dashed lines mark the mean read length; grey lines mark the read-length bounds of the read filter (see Supplementary Table 3 for the filters on read lengths for each library).

**Supplementary Fig. 3.**
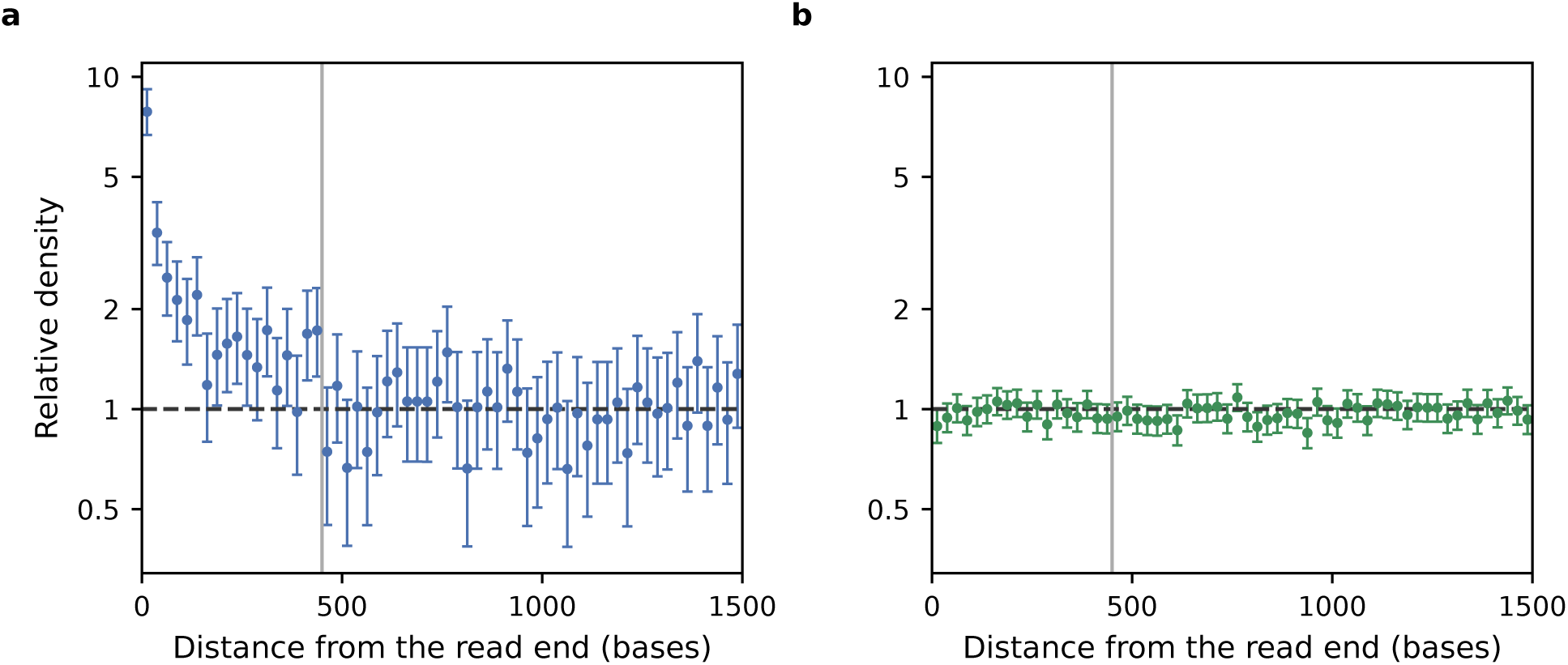
Relative density of singletons and control variants as a function of the distance from the end of the read. Variants called in the untrimmed reads of the eight libraries, at sites that passed the mask and the coverage filter. a, Relative density of singletons. b, Relative density of non-reference alleles found on two or more molecules of the library. Distance is from the nearest end of the forward strand consensus read, in 25-base bins, for reads of at least 3 kb; callable bp within 5 bases of the read end are not counted. The density in each bin is shown relative to the density in the read interior (i.e., sites more than 1,500 bases from both ends of reads of at least 4 kb), which we estimated separately for each library. Error bars are exact 95% Poisson confidence intervals. Grey lines mark the 450 bases trimmed from each read end (see Methods).

**Supplementary Fig. 4.**
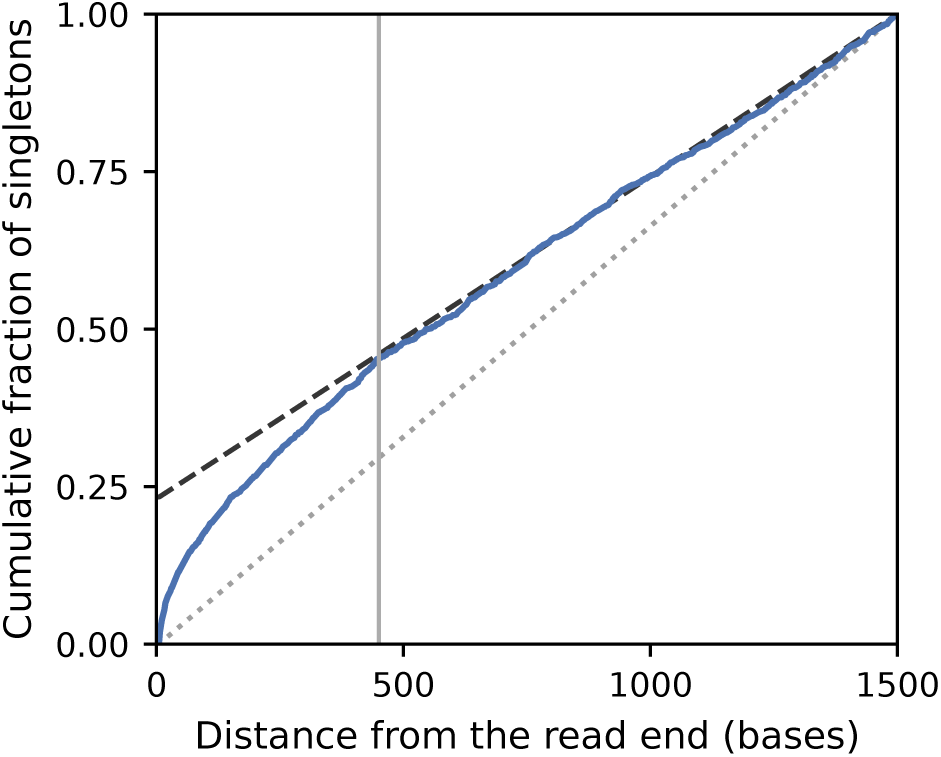
Cumulative fraction of singletons as a function of the distance from the end of the read. The blue curve is the cumulative fraction of singletons within 1,500 bases of either end of the forward strand consensus read. The dotted line is the cumulative fraction of singletons expected in the absence of edge errors, i.e., reflective only of callable bps. The dashed line is the expectation if all edge errors were to occur at the first position in the read. We trimmed 450 bases (grey line), beyond which the two lines converge, and estimated the false discovery rate to be 0.6% (one-sided 95% upper confidence bound 5.4%; see Methods).

**Supplementary Fig. 5.**
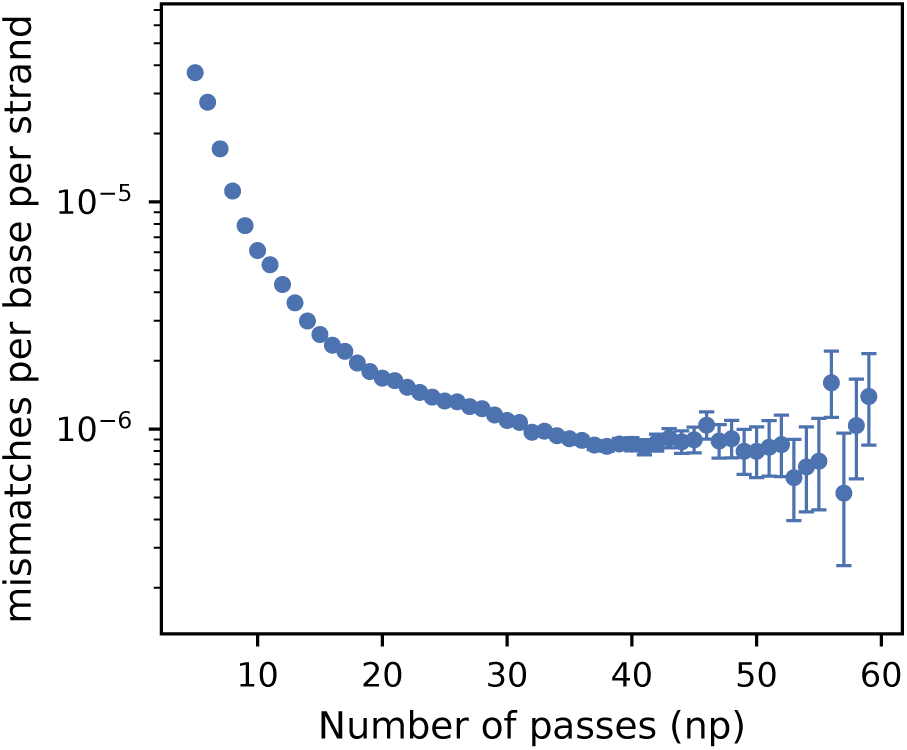
Mismatch rate as a function of the number of passes. Mismatches per base per strand by the number of passes of the strand consensus read (np). Error bars are exact 95% Poisson confidence intervals.

**Supplementary Fig. 6.**
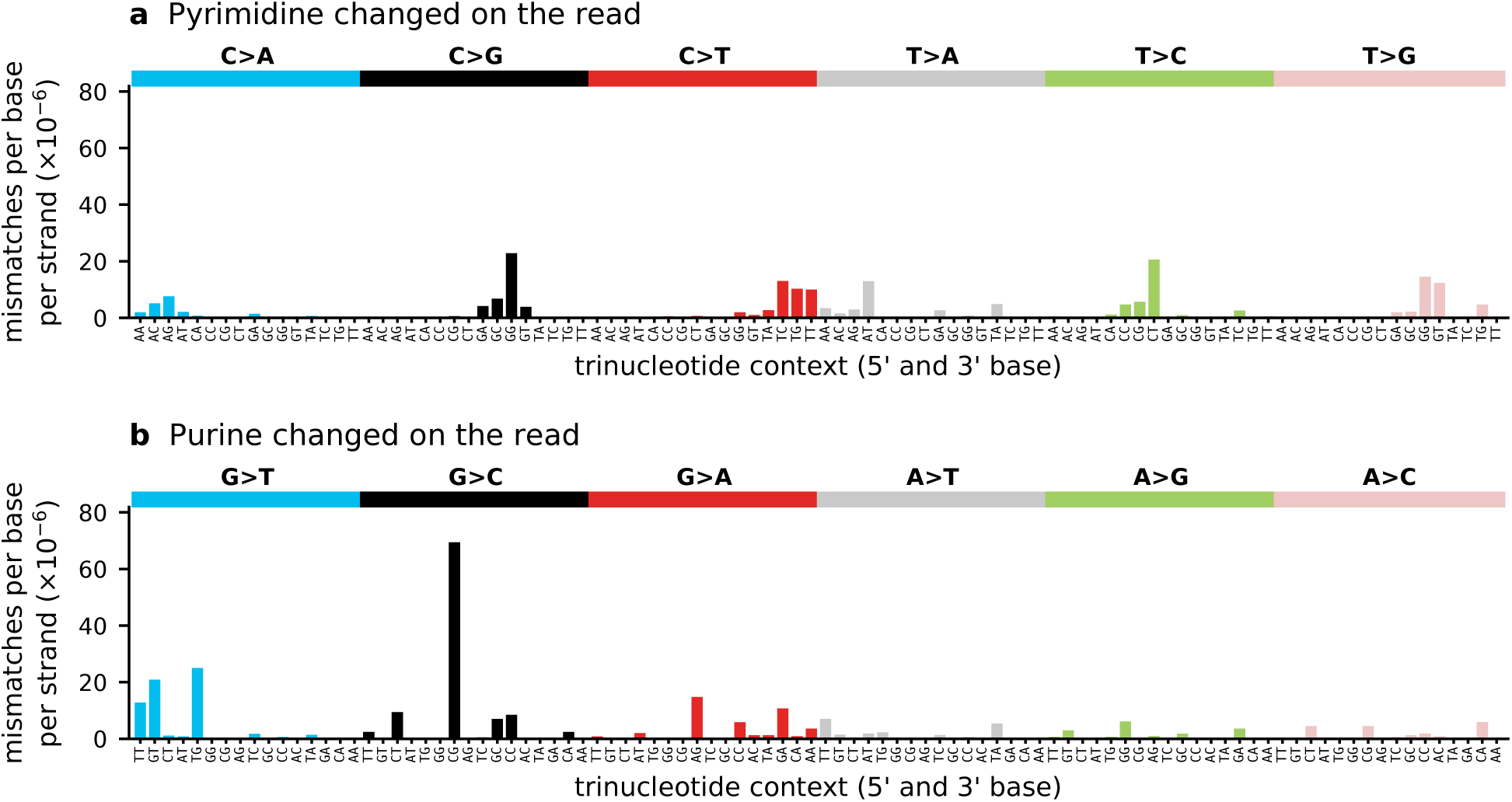
Mismatch rates of each substitution and its reverse complement. a, Rate of each substitution in which a pyrimidine is changed on the read, as the read shows it 5*^′^ →* 3*^′^*. b, Rate of the reverse-complementary substitution, in which a purine is changed on the read, drawn under the corresponding bar of a. Rates are mismatches per base per strand over all strand consensus reads, each divided by its number of opportunities (bases at which the read have the same trinucleotides). All classes of passes are pooled here.

**Supplementary Fig. 7.**
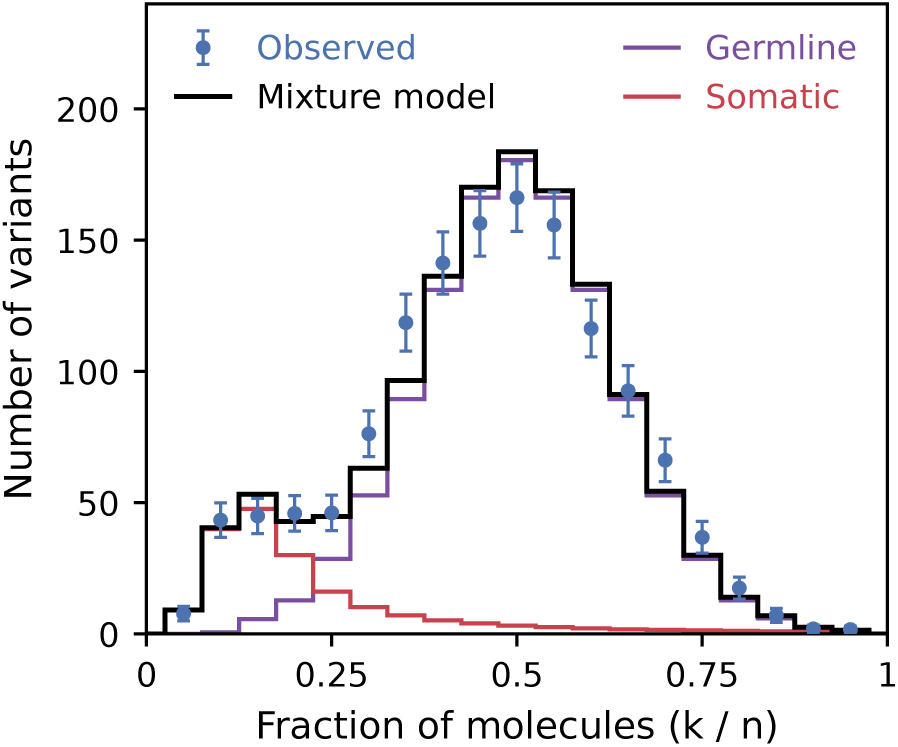
Fraction of molecules carrying the non-reference allele and the mixture model fit. Distribution of the fraction *k/n* of molecules covering the site that carry the non-reference allele, for variants carried by at least two of *n* molecules, 2 *≤ k ≤ n −* 1. Shown are the 1,343 variants across 7 animals (for the 60-week-old mouse, *k* and *n* were summed over its two libraries). Blue dots are the observed counts for each 0.05 bin of *k/n*, with Poisson standard errors; binning is for display purpose only. The data were fitted by maximum likelihood with a mixture model (black line) with two components: germline variants (86%, purple line), for which *k* follows a binomial distribution with mean *n/*2, and somatic mutations (14%, red line), for which *k* follows the distribution expected under a pure-birth process (see Methods).

**Supplementary Fig. 8.**
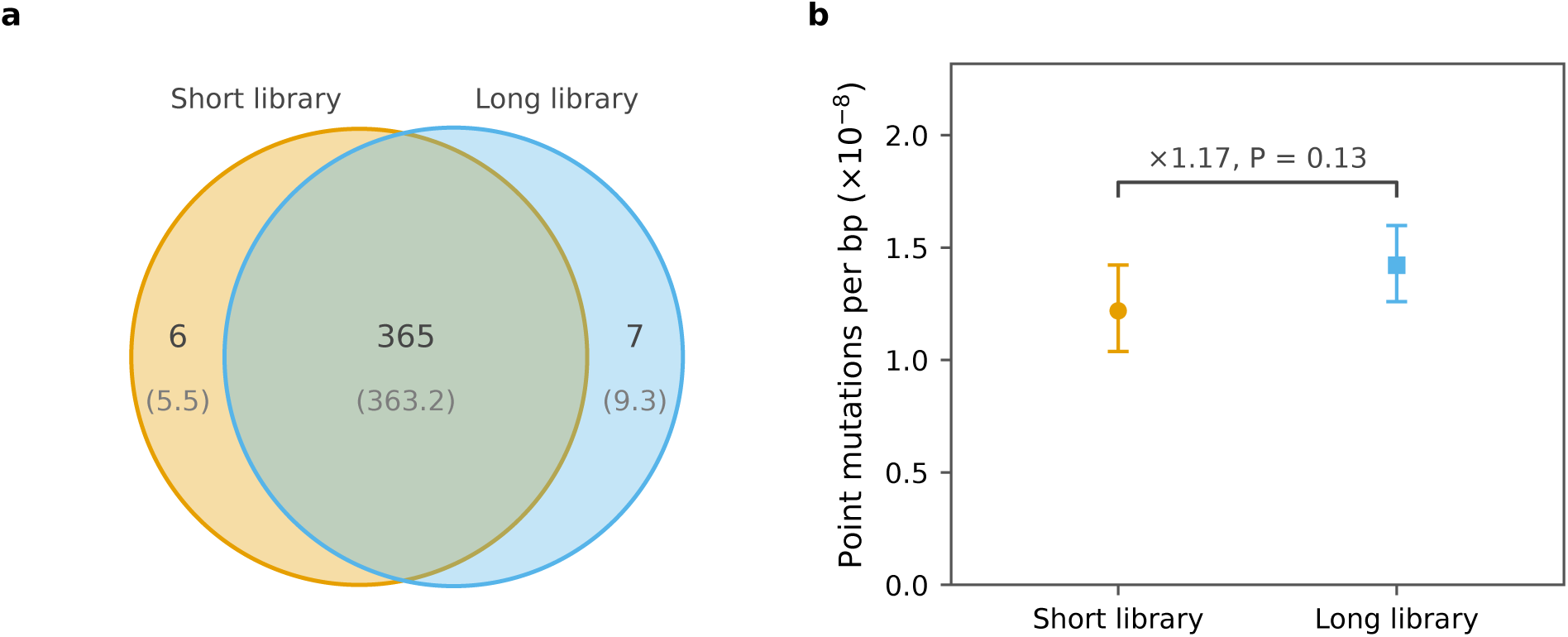
Concordance of the short and long libraries of the 60-week-old mouse. Both panels use the sites callable in both libraries. a, Venn diagram of variants carried by at least two molecules in either library, with the number expected by chance in parentheses. b, Somatic point mutations per bp of each library. Error bars are exact 95% Poisson confidence intervals, and the bracket reports the ratio of the long to the short library and the *P* value of the exact conditional test for two Poisson rates.

**Supplementary Fig. 9.**
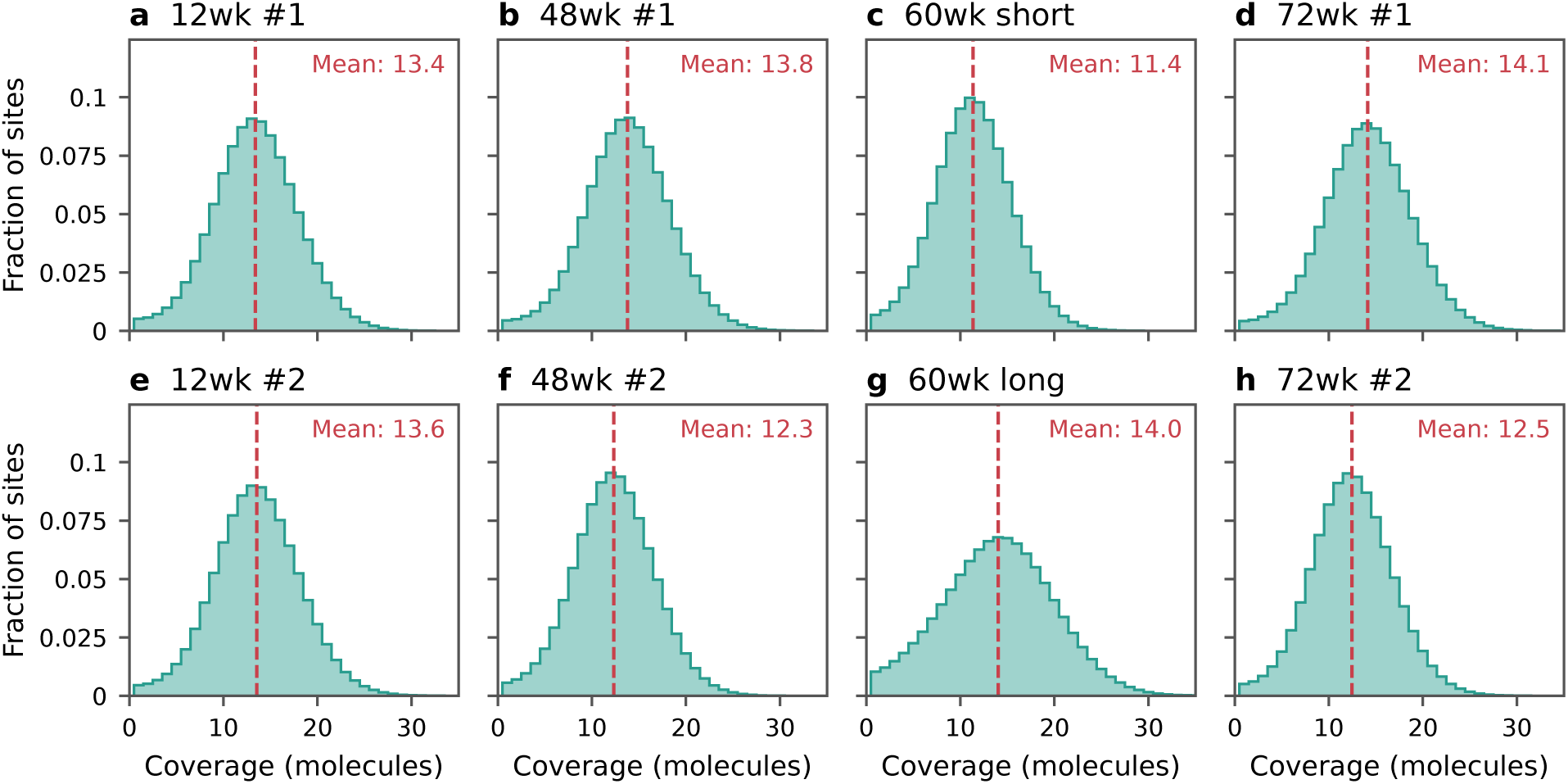
Distribution of untrimmed coverage in each library. Only autosomal sites covered by at least one molecule are included (on average 90.7% of the autosomal genome, Supplementary Table 4). Dashed lines mark the mean coverage across covered sites.

**Supplementary Fig. 10.**
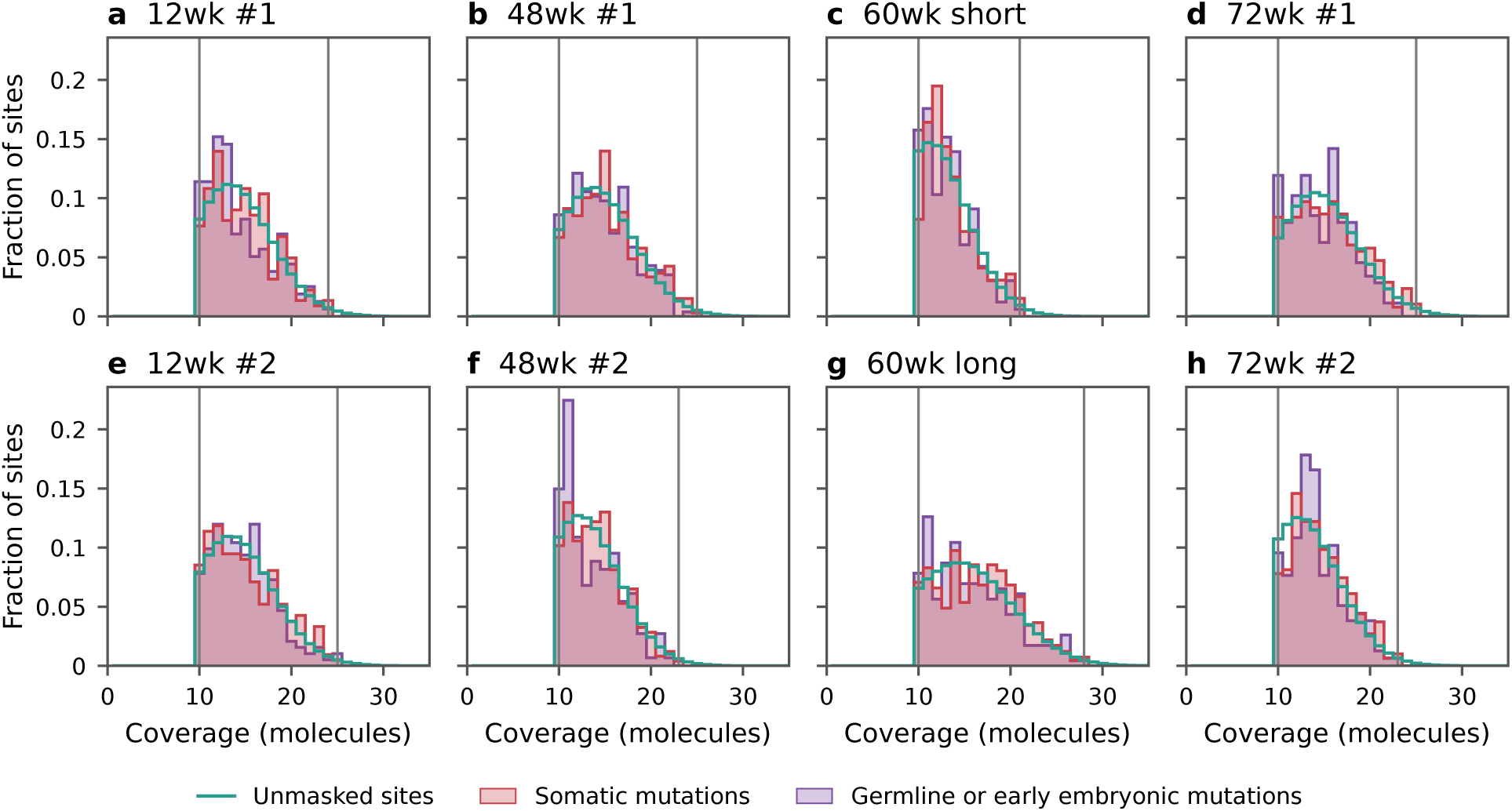
Untrimmed coverage at the somatic mutations and germline or early embryonic mutations of each library. Distribution of untrimmed coverage at the somatic mutations (red), at the germline or early embryonic mutations (purple) and at all unmasked sites with an untrimmed coverage of at least 10 (teal line), each as a fraction of its own sites. Grey lines mark the minimum (10 molecules) and the 99th %-tile coverage of the coverage filter.

**Supplementary Fig. 11.**
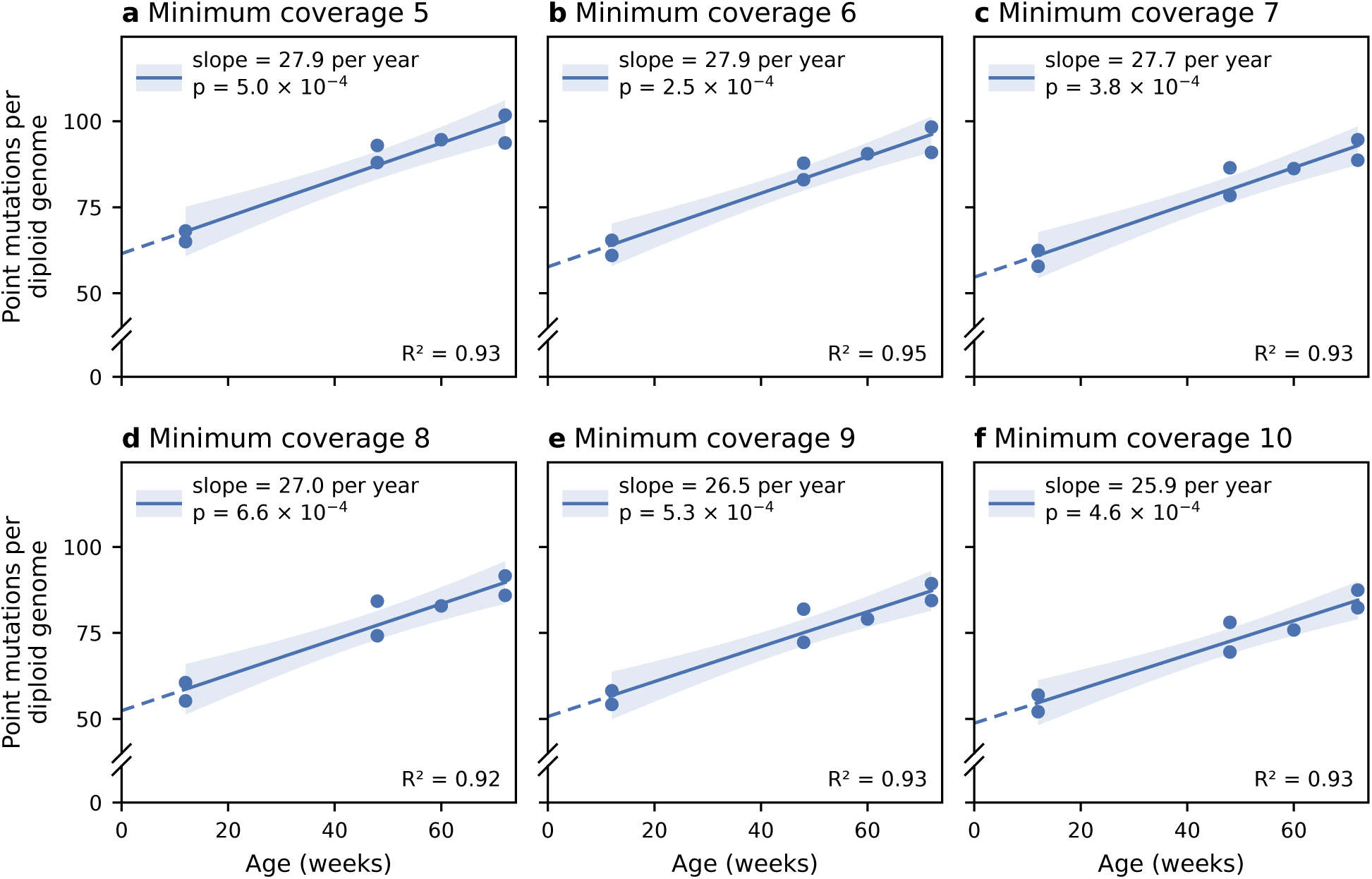
Point mutations per diploid genome as a function of age, for minimum coverages of 5 to 10 molecules. a–f, Minimum coverage 5 (a) to 10 (f). The minimum coverage applies to both the somatic mutations and the callable bps. As it rises from 5 to 10, the total number of point mutations falls from 3,088 to 2,289. Blue dots are the seven animals (the two libraries of the 60-week-old mouse pooled), shown as the number of point mutations per bp multiplied by the diploid genome size (5.28 Gb). The line is the ordinary least squares regression fitted to the seven animals, dashed from the youngest animals back to age 0, where it meets the y axis at the intercept. The shaded area is the 95% confidence interval of the regression line. Each legend gives the slope (point mutations per diploid genome per year) and the *P* -value (two-sided *t* test); *R*^2^ is shown in the lower right of each panel. Panel f is the fit shown in Fig. 2.

**Supplementary Fig. 12.**
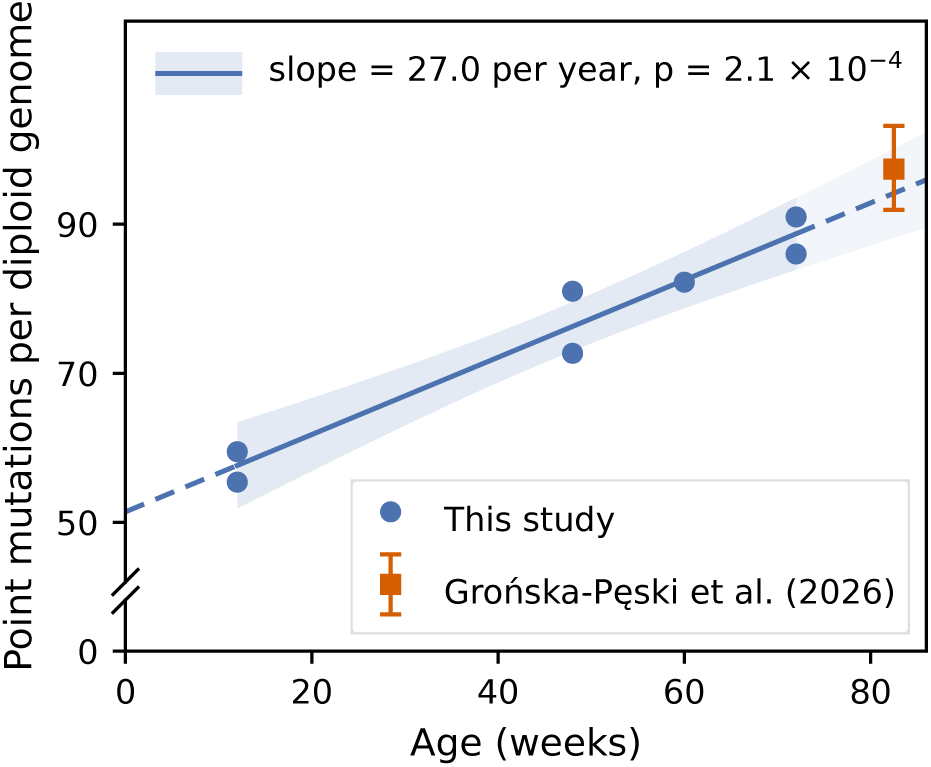
**Somatic point mutations as a function of age, compared with Grońska-Pęski et al.**^19^ . Blue dots are the seven animals (the two libraries of the 60-week-old mouse pooled). Following their approach, here, we corrected for trinucleotide composition: for each mouse, the point mutation rate in each of the 32 trinucleotide contexts is weighted by the mutational opportunities on the autosomes. The orange square and whiskers denote the estimate for cerebellar neurons of 82.5-week-old mice fed *ad libitum*, and its 95% confidence interval (Grońska-Pęski et al.^19^ , their Supplementary Table 6). In that study, the correction for trinucleotide context was based on the GRCm38 genome and extrapolated to a diploid genome with the same genome size as in our case. The line is from an ordinary least squares regression fitted to the seven animals (slope: 27.0 point mutations per diploid genome per year; 95% CI: 19.8 to 34.2; *P* = 2.1 *×* 10*^−^*^4^, two-sided *t* test; *R*^2^ = 0.95). The shaded area is the 95% confidence interval of the regression line, and is lighter where it is extrapolated.

**Supplementary Fig. 13.**
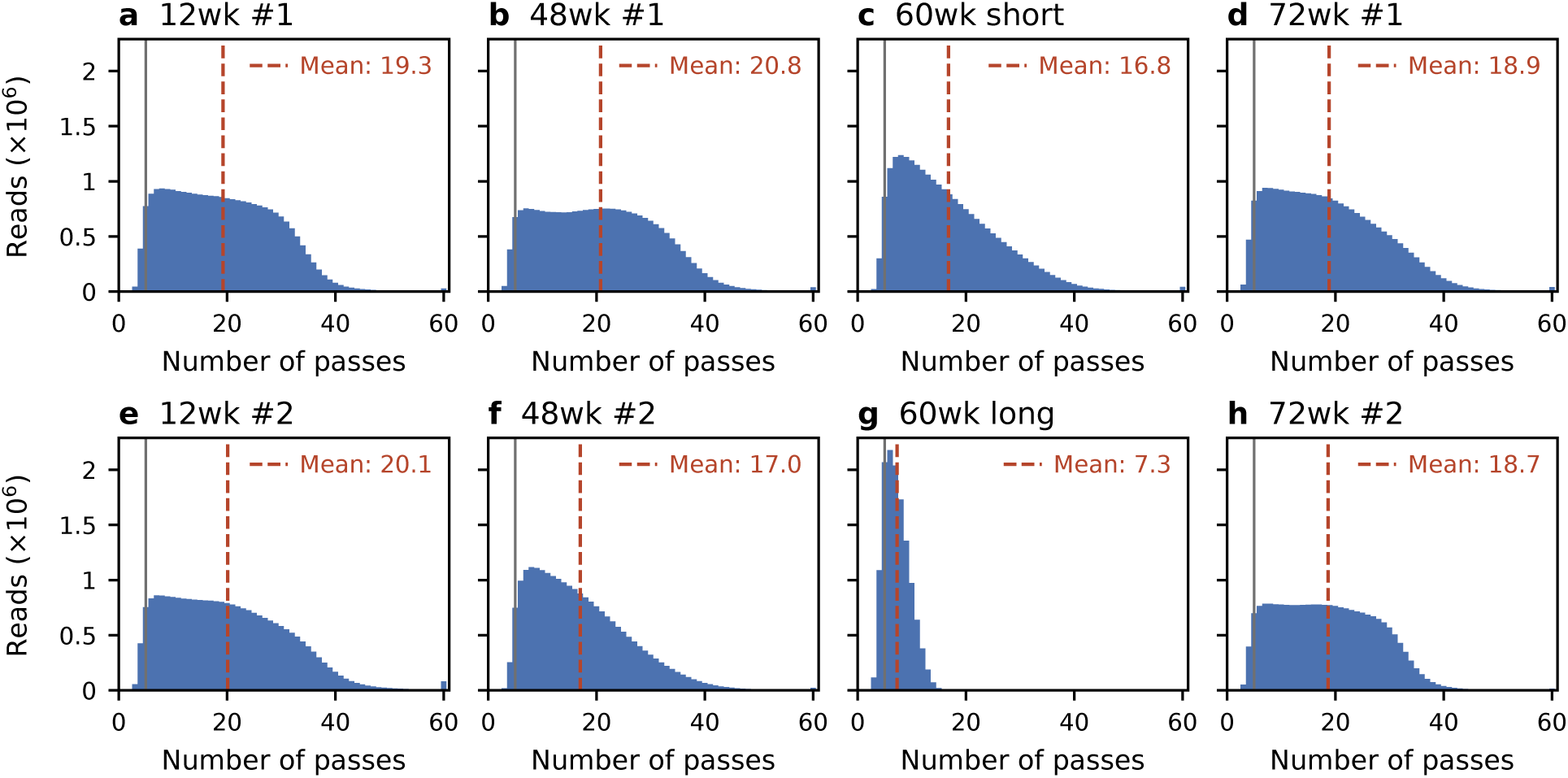
Distribution of number of passes of per strand consensus reads in each library. Dashed lines mark the mean; grey lines mark the minimum of five passes of the read filter.

**Supplementary Fig. 14.**
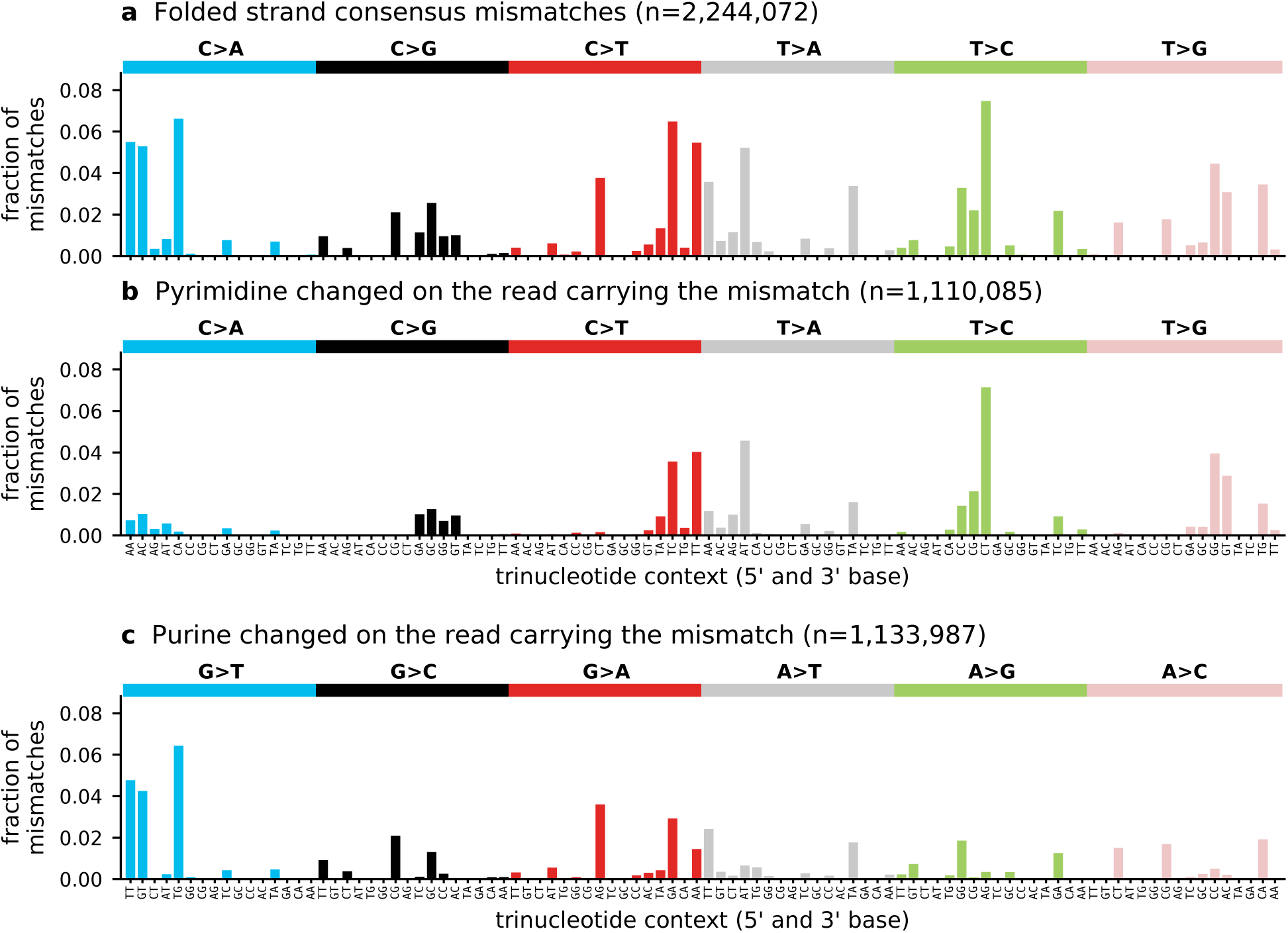
Spectrum of strand consensus mismatches. a, All mismatches, folded to the six pyrimidine substitution classes. b, Mismatches in which a pyrimidine was changed on the divergent read, as the read shows them 5*^′^ →* 3*^′^*. c, Mismatches in which a purine was changed on the divergent read, as the read shows them 5*^′^ →* 3*^′^*, each drawn under the channel of the complementary pyrimidine change. Bars are fractions of all mismatches, so that mismatches in b and c add up to those shown in a. *n* is the number of mismatches.

**Supplementary Fig. 15.**
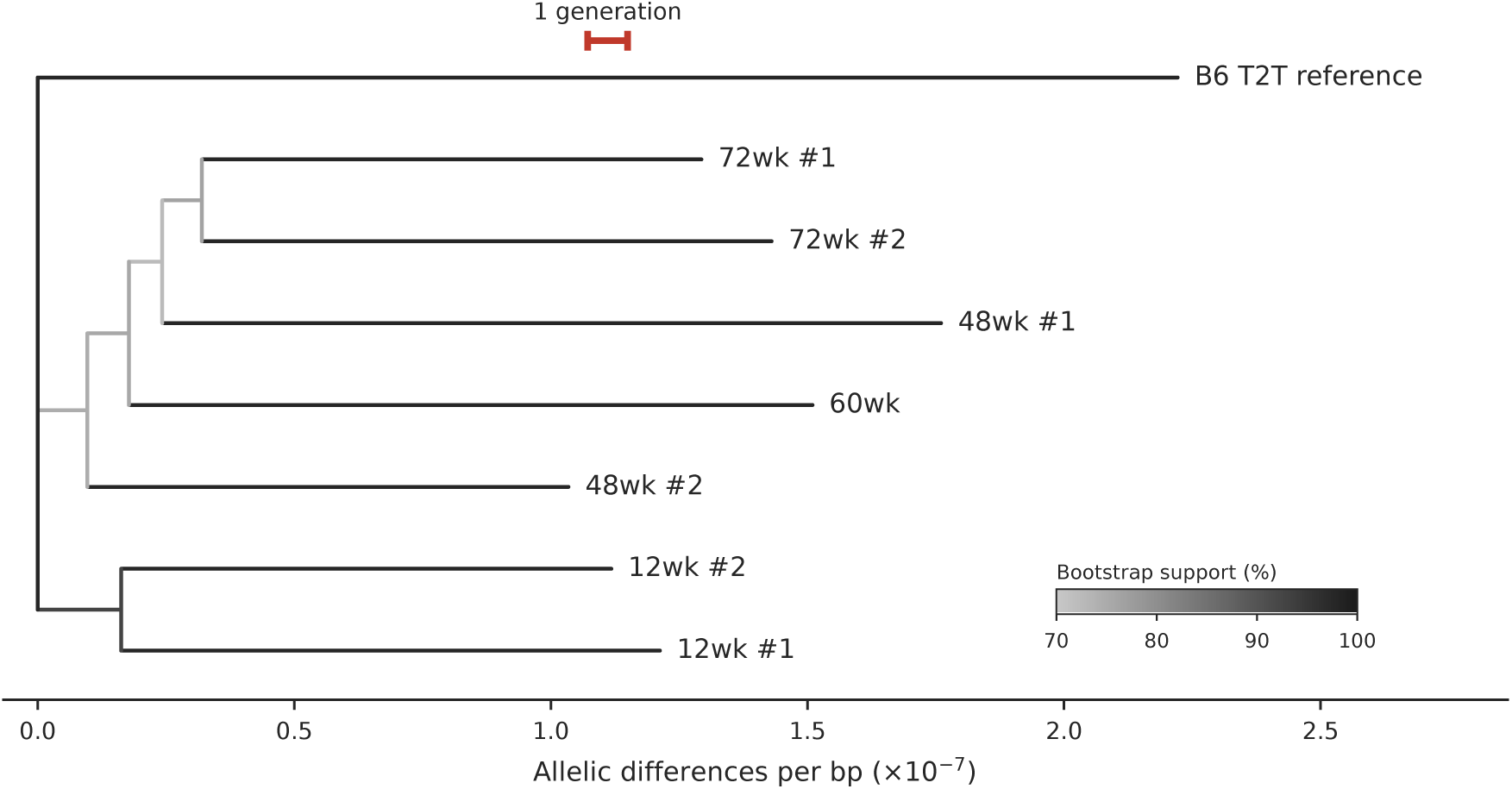
Neighbour-joining tree of the seven animals. The tree is built from the genotypes of the animals at 501 sites with a non-reference allele carried by at least two molecules in a mouse (Supplementary Table 7); to this end, the two libraries of the 60-week-old mouse were pooled. The reference genome was used as an outgroup (drawn at the top). Branch lengths are allelic differences per bp, and branches are shaded by bootstrap support (10,000 replicates). The red scale bar corresponds to the germline mutations that a mouse inherits in one generation, based on a cross of C57BL/6 and 129S5 mice ^38^, i.e., *∼*5 mutations in the 0.63 Gb used to build the tree.

## Notes

### Competing Interest Statement

The authors have declared no competing interest.

